# Functional connectivity fingerprints of the frontal eye fields and inferior frontal junction in the dorsal vs. ventral prefrontal cortex

**DOI:** 10.1101/2022.06.04.494797

**Authors:** Orhan Soyuhos, Daniel Baldauf

## Abstract

Neuroimaging evidence suggests that the frontal eye field (FEF) and inferior frontal junction (IFJ) govern the encoding of spatial and non-spatial (such as feature- or object-based) representations, respectively, both during visual attention and working memory tasks. However, it is still unclear whether such contrasting functional segregation is also reflected in their underlying functional connectivity patterns. Here, we hypothesized that FEF has predominant functional coupling with spatiotopically organized regions in the dorsal (‘where’) visual stream, whereas IFJ has predominant functional connectivity with the ventral (‘what’) visual stream. We applied seed-based functional connectivity analyses to temporally high-resolving resting-state magnetoencephalography (MEG) recordings. We parcellated the brain according to the multimodal Glasser atlas and tested, for various frequency bands, whether the spontaneous activity of each parcel in the ventral and dorsal visual pathway has predominant functional connectivity with FEF or IFJ. The results show that FEF has a robust power correlation with the dorsal visual pathway in beta and gamma bands. In contrast, anterior IFJ (IFJa) has a strong power coupling with the ventral visual stream in delta, beta, and gamma oscillations. Moreover, while FEF is directly phase-coupled with the superior parietal lobe in the beta band, IFJa is directly phase-coupled with the middle and inferior temporal cortex in delta and gamma oscillations. We argue that these intrinsic connectivity fingerprints are congruent with each brain region’s function. Therefore, we conclude that FEF and IFJ have dissociable connectivity patterns that fit their respective functional roles in spatial vs. non-spatial top-down attention and working memory control.

## 1 INTRODUCTION

At any moment, we face an overwhelming stream of rich information from the surrounding environment. Our brain must continuously re-evaluate all the incoming sensory stimulation to prioritize what is relevant for our behavioral goals (Baldauf & Deubel, 2010; Desimone & Duncan, 1995; Fecteau & Munoz, 2006). The prefrontal cortex (PFC), the maestro in orchestrating a range of cognitive functions in the human brain, plays an essential role in this respect (Duncan, 2001; Fuster, 2001; Miller & Cohen, 2001; Rossi et al., 2008). It crucially contributes to the central functions of selective attention and working memory by top-down modulating priority for incoming stimuli according to the context of the ongoing task demands (Jerde et al., 2012; Ptak, 2012). Its distinct control circuits’ interareal communications with the rest of the cortex enable the brain to focus its limited perceptual resources on specific locations in space, or on certain features or objects (Baldauf & Desimone, 2014; Bichot et al., 2019; Corbetta & Shulman, 2002; Gregoriou et al., 2009; Gross et al., 2004; Mangun, 1995; Siegel et al., 2008). In this context, human neuroimaging techniques and invasive recordings in primates suggest that the representational content encoded during visual selective attention and working memory can further segregate the functional specialization within PFC (Carrasco, 2011; Cavada & Goldman-Rakic, 1989; Giesbrecht et al., 2003; Goldman-Rakic, 1988; McCarthy et al., 1996; Scalaidhe et al., 1999; Paneri & Gregoriou, 2017; Romanski, 2004; Wilson et al., 1993). In particular, two specific control sites located in the posterior lateral PFC (plPFC), namely the Frontal Eye Field (FEF) (Kelley et al., 2008; Petit & Pouget, 2019; Vernet et al., 2014) and the Inferior Frontal Junction (IFJ) (Baldauf & Desimone, 2014; Brass et al., 2005; Derrfuss et al., 2005; Muhle-Karbe et al., 2014), show differential neural activity depending on the stimulus and task domain.

Experiments in which attention is covertly directed to a visuospatial cue have led to increased neural activity in FEF, apart from its oculomotor function (Armstrong et al., 2009; Thompson et al., 2005; Thompson & Bichot, 2005). Additionally, muscimol injection and suprathreshold microstimulation techniques revealed FEF’s crucial role in spatial attention (Moore & Armstrong, 2003; Moore & Fallah, 2004; Wardak et al., 2006). In this respect, Thompson and Bichot (2005) reported the existence of a visual salience map in primate FEF and noted that its initial neural activities are not selective for non-spatial features such as color or shape, but that their activity evolves over time to select the target location. Along these lines, human neuroimaging and monkey studies indicated that FEF exerts top-down regulatory influences on the extrastriate visual cortex and increases the response gain to retinotopically corresponding targets directly or indirectly through its connections to the intraparietal sulcus (IPS) (Bressler et al., 2008; Gregoriou et al., 2009, 2012; Kelley et al., 2008; Merrikhi et al., 2017). Moreover, transcranial magnetic stimulation (TMS) studies advanced these correlational findings with causal evidence and pointed out FEF’s key function in the top-down control of visuospatial attention (Heinen et al., 2017; Marshall et al., 2015; Veniero et al., 2021).

In contrast, experimental designs that require attention to non-spatial features or object representations have consistently invoked neural responses in the inferior frontal cortex around IFJ. This evidence mainly accumulated from human neuroimaging studies due to the lack of corresponding sulcal landmarks in the nonhuman primate brain (Bedini & Baldauf, 2021; Donahue et al., 2018). Nevertheless, the studies on the ventral pre-arcuate of monkey PFC (VPA; regarded as the homolog of IFJ) have provided important insights into the region’s functioning. For instance, Bichot and colleagues (2015) reported VPA as the source of feature selection. They showed that the inactivation of VPA reduces the feature-based modulation in ipsilateral V4 and impairs monkeys’ performance in detecting target objects in the contralateral visual field (Bichot et al., 2019). Likewise, human neuroimaging and TMS studies reported a top-down modulatory role of IFJ in object- and feature-based attention and working memory (Baldauf & Desimone, 2014; Meyyappan et al., 2021; Zanto et al., 2010, 2011; Zhang et al., 2018). For instance, Zanto and colleagues (2010) found that the functional coupling between the right IFJ and V4 was predictive of the extent of attentional modulation during attention to color. Additionally, they demonstrated that the perturbation of IFJ weakens working memory performance for color features (Zanto et al., 2011). Following this, through directional phase coupling, Baldauf and Desimone (2014) additionally showed that IFJ is the source of attentional modulation of object representations in the inferior temporal (IT) cortex and the driver of non-spatial, object-based attention.

Overall, the evidence from task-based studies suggests that the encoding of to-be-attended or to-be-remembered representational contents functionally segregates PFC and reveals two distinct control circuits in the dorsal and ventral PFC: FEF and IFJ. While FEF is involved in the top-down control of spatial attention and working memory, IFJ contributes to the top-down control of non-spatial (feature- and object-based) attention and working memory (for a review see: Bedini & Baldauf, 2021). Nonetheless, it is still unclear to what extent this functionally specific contrast is reflected in these regions’ underlying functional connectivity profiles.

### 1.1 Research question and hypothesis

The current study aimed to analyze the intrinsic functional connectivity fingerprints of FEF and IFJ to support their functional segregation within PFC. We argue that FEF and IFJ’s interareal communication with regions in the dorsal (‘where’) and ventral (‘what’) visual pathways (Goodale & Milner, 1992; Mishkin et al., 1983) would be a prerequisite for their functional involvement in spatial and non-spatial attention and working memory control, respectively; similarly to the structural connectivity fingerprints of areas in IT cortex being predictive of their functional selectivity (Osher et al., 2016; Saygin et al., 2011). In this respect, we hypothesize that FEF has predominant functional connectivity with spatiotopically organized areas in the dorsal visual stream, whereas IFJ has predominant functional connectivity with feature- or object-encoding areas in the ventral visual pathway. To this end, we performed a region of interest (ROI) analysis and explored FEF and IFJ’s functional connectivity to the rest of the brain. Additionally, we inquired about the dominant directionality of these interactions between FEF/IFJ and the visual streams. To this aim, we used various functional connectivity metrics (both phase- and power-based) and epoch lengths to consider their effects on the consistency of results.

## 2 MATERIALS AND METHODS

### 2.1 Participants

We benefited from the 1200 Subjects Release of Human Connectome Project (HCP) dataset that includes resting-state MEG (rMEG) recordings and structural MRI of 95 healthy participants aged between 22 and 35 (Larson-Prior et al., 2013; Van Essen et al., 2013). The Washington University institutional review board approved data acquisition protocols for HCP. Each subject was asked for verbal consent before the sessions. Thirty-six participants are reported to be monozygotic and dizygotic twins. Out of 95 subjects, we selected 55 independent ones (26 female) after randomly excluding one of the pairs from each twin and checking the data quality. The anonymized dataset is provided in ConnectomeDB (db.humanconnectome.org; Hodge et al., 2016).

### 2.2 Procedure

Subjects typically spent 2 or 3 days completing the HCP protocol that includes behavioral tests, resting-state functional magnetic resonance imaging (MRI), task-evoked functional MRI, diffusion-weighted MRI, and task-evoked MEG in addition to structural MRI and rMEG sessions. MEG recordings always took place before MRI sessions to prevent the potential effect of residual magnetization caused by the MRI scanner on the superconducting MEG sensors (Van Essen et al., 2013).

### 2.3 MRI data acquisition and preprocessing

Washington University - University of Minnesota Consortium of the Human Connectome Project (WU-Minn HCP Consortium) performed Structural MRI on a customized Siemens 3T “Connectome Skyra” at Washington University (Uğurbil et al., 2013). They acquired two high-resolution T1- (T1w) and T2-weighted (T2w) anatomical images of participants using a standard 32-channel Siemens receive head coil (T1w; 3D MPRAGE; 0.7 mm isotropic; FOV, 224 × 224; 256 sagittal slices; TR=2400 ms; TE=2.14 ms; inversion time (TI), 1000 ms; 8-degree flip angle) (T2w; 3D T2-SPACE; 0.7 mm isotropic; FOV, 224 × 224; 256 sagittal slices; TR=3200 ms; TE=565 ms; variable flip angle). Experts reviewed the quality of scans, considering the tissue contrast, blurriness, and artifacts. If necessary additional scans were arranged for a second time. Also, a radiologist examined the existence of any neuroanatomical anomalies. While the normal variations were noted, subjects with abnormal brain anatomy were excluded from the dataset.

ConnectomeDB provides preprocessed structural data resulting from the HCP structural pipeline (Glasser et al., 2013). The second part of this pipeline is mainly based on a modified version of FreeSurfer’s recon-all script to reconstruct a two-dimensional cortical surface from a three-dimensional volume (Dale et al., 1999; Fischl, Sereno, & Dale, 1999). We used the resulting files provided inside the Structural Extended Preprocessed package in the ConnectomeDB for the source reconstruction of MEG data.

### 2.4 MEG data acquisition

The rMEG recordings were acquired on a whole head Magnes 3600 scanner (4D Neuroimaging, San Diego, CA) in a magnetically shielded room at Saint Louis University (Larson-Prior et al., 2013). The MEG scanner had 248 magnetometers and 23 reference channels (18 magnetometers and five first-order gradiometers). The data were recorded with a sampling rate of 2034.5101 Hz, delta encoded, and saved in a 4D file format. Electrooculography (EOG), electrocardiography (ECG), and electromyography (EMG) signals were also measured simultaneously to enable offline artifact rejection. Moreover, the experimenters used five Head Position Indicators (HPI) to track the head position in reference to the MEG sensors and marked the anatomical landmarks of nasion and the preauricular points to coregister the SQUID array to each individual’s structural MRI. They used a Polhemus Fastrak-III system for the spatial digitization of localizer coils and fiducial points. Additional points defining the subject’s head shape (about 2400 points) were also digitized to fine-tune the co-registration.

During the rMEG scanning, subjects lay in a supine position. They were instructed to keep their heads still while fixating a red crosshair at the center of the projection. Their head positions were recorded before and after every three consecutive sessions. Each run took approximately 6 minutes. Before rMEG, the empty room and participant noise recordings were also acquired to monitor the sensor noise and the noise stemming from the participant due to residual magnetization effects.

### 2.5 MEG data preprocessing

The ASCII text files that annotate the indexes of bad channels, bad segments, and non-brain components are provided inside the Preprocessed Resting-State package in the ConnectomeDB. To ensure the reproducibility of our results, we decided to utilize them for preprocessing rMEG data. First, the noisy channels and the bad data segments were marked following a visual inspection. Later, the correlation between signals from adjacent sensors and the ratio of their variance to their neighbors were calculated to mark the outliers as bad channels. Additionally, z-scores for all time points in each channel were computed to identify the segments whose score was higher than a given threshold. The clipping and muscle-related artifacts were detected using ft_artifact_clip.m and ft_artifact_muscle.m functions from the FieldTrip toolbox (Oostenveld et al., 2010). Furthermore, an iterative Independent Component Analysis (ICA) was performed to detect faulty channels and bad segments in the raw data by using the list of bad channels and segments identified beforehand as initial input. In subsequent iterations of this ICA, the input was updated by the bad sensors or segments found in the previous iteration until the ICA algorithm could no longer identify any further bad channels/segments.

In the next step of the MEG preprocessing pipeline, ICA was used this time to remove the artifacts related to eye movements, blinks, and heartbeats from the signal. To this end, first, a bandpass (1.3-150 Hz) and a notch Butterworth (59-61 and 119-121 Kz) filter were applied to the MEG, ECG, and EOG data. The previously identified bad channels and segments were cut out from these three recordings. Afterward, ICA was implemented on the preprocessed MEG data twenty times based on different seed values. Power spectral densities (PSD) and power time courses (PTC) were calculated for the independent components (ICs) of each repetition and corresponding time series of ECG and EOG channels. An automatic classification algorithm detected artifacts based on the correlation between ICs and ECG/EOG recordings. It classified ICs into the brain and non-brain components while also considering pink noise and the noise stemming from the power supply (60 and 120 Hz). The classifier’s performance was later confirmed by visual inspection. Ultimately, the best iteration with the highest ratio of brain to artifact components from twenty repetitions was selected. The indices of its brain and non-brain components were reported in ASCII text files. After the removal of the bad channels, bad segments, and artifacts from the rMEG data, we parsed the continuous times series into non-overlapping epochs of either 2 s, 5 s, or 10 s to consider the epoch length’s effect on the connectivity measures (Brookes et al., 2011; Fraschini et al., 2016).

### 2.6 Source reconstruction of MEG signals

For each epoch length, we utilized the Brainstorm toolbox (Tadel et al., 2011) to reconstruct source-level activation on the native cortex of each subject. We first corrected the DC offset of preprocessed rMEG data and downsampled it to 300 Hz. To estimate sensor noise, we computed the noise covariance matrix from raw empty room recording after removing its DC bias. Additionally, we calculated the lead field matrix for an evenly distributed grid of 15002 cortical sources by fitting a single sphere under each sensor (Huang et al., 1999). Based on these gain and noise covariance matrices, we applied the minimum-norm estimation (MNE) method to inverse-model activations in source space (Baillet et al., 2001; Dale et al., 2000). Here, we chose MNE rather than beamformers because it was shown to perform better for spatially distant sources (Hincapié et al., 2017). Afterward, the source level activity for each subject was projected to an MNI average brain template (FreeSurfer fsaverage; Fischl et al., 1999) and parcellated by the multimodal parcellation atlas (MMP1; Glasser et al., 2016). We calculated the scout activations for 360 regions by taking the maximum absolute value across all vertices per scout and multiplying with its sign. We saved them as Fieldtrip structures to perform functional connectivity analyses in the next step.

The MEG Anatomy package provided in the ConnectomeDB enables the automatic co-registration of MEG sensors with the structural MRI. However, it does not allow projecting sources to a standard brain (such as fsaverage). Since we aimed to interpolate source-level activation of each subject onto the fsaverage template for the group level analysis, we used the pial surface provided in the Structural Extended Preprocessed package to reconstruct sources. In this way, we could also increase the resolution of the cortex from 8004 to 15002 vertices for a better representation of cortical folds. However, we still had to use the MEG Anatomy package to retrieve the coordinates of fiducial points for each subject since the individual MRIs were defaced, and the fiducial points were not stated explicitly.

### 2.7 Functional connectivity and directionality analyses

In a second step, we conducted seed-based functional connectivity analyses with seed regions of FEF, anterior IFJ (IFJa), and posterior IFJ (IFJp) in both hemispheres by employing the FieldTrip toolbox (Oostenveld et al., 2010). We first carried out frequency analyses on time series data over all epochs while using the multiple tapers based on the Slepian sequence. The frequencies of interest were the delta (1-4 Hz), theta (4-8 Hz), alpha (8-13 Hz), beta (13-30 Hz), and gamma (30-100 Hz) band with the number of spectral smoothing boxes of 1, 1, 1, 2 and 10, respectively. We computed complex Fourier-spectra between all brain parcels for each frequency band. Based on this Fourier representation of data, we calculated the functional connectivity between each of the six seed regions and the rest of the 359 brain parcels. We decided to use both phase- and amplitude-based functional connectivity measures to take into account the possible distinct roles of phase- and amplitude-coupling (Daffertshofer et al., 2018; Siems & Siegel, 2020). In particular, we chose the imaginary part of the Coherency (iCOH) (Nolte et al., 2004), debiased weighted Phase Lag Index (dwPLI) (Vinck et al., 2011), and orthogonalized Power Envelope Correlation (oPEC) (Hipp et al., 2012) since these metrics are corrected for spatial leakage and have higher group-level repeatability compared to other methods (Bastos & Schoffelen, 2016; Brookes et al., 2011; Colclough et al., 2016; Duan et al., 2021).

We performed specific region-of-interest (ROI) and whole-brain exploratory analyses based on the connectivity metrics mentioned above. For ROI analyses, the target MMP1 parcels in Table 1 were chosen based on a systematic bibliographical search of the relevant literature (Felleman & Van Essen, 1991; Glasser et al., 2016; Goodale & Milner, 1992; Kravitz et al., 2011, 2013; Mishkin et al., 1983) and the list reported in Zimmermann et al. (2018). We first selected the mid-higher visual areas that belong to the dorsal or ventral visual pathway. Additionally, we used the retinotopy atlas by Wang et al. (2015) to identify putative retinotopic parcels of the MMP1 atlas and include them in our definition of the dorsal visual stream, as the topographic organization is generally considered a defining feature of this stream. We excluded parcels near the medial wall, parcels from the superior temporal lobe, and the temporo-parietal junction since they seem to be characterized by distinctive connectional profiles (Kravitz et al., 2011, 2013; Mars et al., 2011). This led to a total of 33 regions with specific visual properties and selectivity to visual features (Table 1). We tested whether the spontaneous activity of each parcel in the ventral and dorsal visual pathway has higher functional connectivity with FEF or IFJ. Following this, we additionally questioned whether the regions that do not show preferential connectivity, neither with FEF nor with IFJ, have a statistically significant intrinsic functional connectivity with any of these seed regions when tested against the mean of whole-brain connectivity values per frequency band.

**TABLE 1.** List of parcels from the MMP1 atlas for ROI analyses. We selected ROIs in the visual cortex as belonging to the ventral or dorsal visual stream by considering previous arguments for their inclusion in the dorsal/ventral pathway, receptive field sizes, and the existence of a topographical organization.

| Parcel | Description | Visual Stream | Topographical Organization |
| --- | --- | --- | --- |
| Dorsal Visual Pathway |  |  |  |
| V6 | Sixth Visual Area | Dorsal (Glasser et al., 2016) | ✓ (Kravitz et al., 2011) |
| V6A | Area V6A | Dorsal (Glasser et al., 2016; Zimmermann et al., 2018) | ✓ (Kravitz et al., 2011; Wang et al., 2015) |
| V7 | Seventh Visual Area | Dorsal (Glasser et al., 2016) | ✓ (Glasser et al., 2016; Wang et al., 2015) |
| IPS1 | IntraParietal Sulcus Area 1 | Dorsal (Glasser et al., 2016; Zimmermann et al., 2018) | ✓ (Wang et al., 2015) |
| IP1 | Area IntraParietal 1 | Dorsal (Zimmermann et al., 2018) | ✓ (Wang et al., 2015) |
| MIP | Medial IntraParietal Area | Dorsal (Zimmermann et al., 2018) | ✓ (Kravitz et al., 2011; Wang et al., 2015) |
| VIP | Ventral IntraParietal Complex | Dorsal (Zimmermann et al., 2018) | ✓ (Felleman & Van Essen, 1991; Kravitz et al., 2011; Wang et al., 2015) |
| LIPd | Area Lateral IntraParietal dorsal | - | ✓ (Felleman & Van Essen, 1991; Kravitz et al., 2011; Wang et al., 2015) |
| LIPv | Area Lateral IntraParietal ventral | Dorsal (Zimmermann et al., 2018) | ✓ (Felleman & Van Essen, 1991; Kravitz et al., 2011; Wang et al., 2015) |
| 7AL | Lateral Area 7A | Dorsal (Zimmermann et al., 2018) | ✓ (Wang et al., 2015) |
| 7PC | Area 7PC | Dorsal (Zimmermann et al., 2018) | ✓ (Wang et al., 2015) |
| 7PL | Lateral Area 7P | Dorsal (Zimmermann et al., 2018) | ✓ (Wang et al., 2015) |
| V3A | Area V3A | Dorsal (Glasser et al., 2016; Goodale & Milner, 1992) | ✓ (Glasser et al., 2016; Felleman & Van Essen, 1991; Kravitz et al., 2013; Wang et al., 2015) |
| V3B | Area V3B | Dorsal (Glasser et al., 2016) | ✓ (Wang et al., 2015) |
| V3CD | Area V3CD | - | ✓ (Wang et al., 2015) |
| LO3 | Area Lateral Occipital 3 | - | ✓ (Wang et al., 2015) |
| MT | Middle Temporal Area | Dorsal (Goodale & Milner, 1992) | ✓ (Glasser et al., 2016; Felleman & Van Essen, 1991; Kravitz et al., 2013) |
| Ventral Visual Pathway |  |  |  |
| V8 | Eighth Visual Area | Ventral (Glasser et al., 2016) | - |
| VMV1 | VentroMedial Visual Area 1 | Ventral (Glasser et al., 2016) | - |
| VMV2 | VentroMedial Visual Area 2 | Ventral (Glasser et al., 2016) | - |
| VMV3 | VentroMedial Visual Area3 | Ventral (Glasser et al., 2016) | - |
| PHT | Area PHT | Ventral (Zimmermann et al., 2018) | - |
| PH | Area PH | Ventral (Zimmermann et al., 2018) | - |
| TE1a | Area TE1 anterior | Ventral (Goodale & Milner, 1992; Kravitz et al., 2011, 2013; Zimmermann et al., 2018) | - |
| TE1m | Area TE1 middle | Ventral (Goodale & Milner, 1992; Kravitz et al., 2011, 2013; Zimmermann et al., 2018) | - |
| TE1p | Area TE1 posterior | Ventral (Goodale & Milner, 1992; Kravitz et al., 2011, 2013; Zimmermann et al., 2018) | - |
| TE2a | Area TE2 anterior | Ventral (Goodale & Milner, 1992; Kravitz et al., 2011, 2013; Zimmermann et al., 2018) | - |
| TE2p | Area TE2 posterior | Ventral (Goodale & Milner, 1992; Kravitz et al., 2011, 2013; Zimmermann et al., 2018) | - |
| TF | Area TF | Ventral (Goodale & Milner, 1992; Kravitz et al., 2011; Zimmermann et al., 2018) | - |
| TGv | Area TG ventral | Ventral (Kravitz et al., 2013; Zimmermann et al., 2018) | - |
| FFC | Fusiform Face Complex | Ventral (Glasser et al., 2016) | - |
| PIT | Posterior Infero Temporal | Ventral (Glasser et al., 2016) | - |
| VVC | Ventral Visual Complex | Ventral (Glasser et al., 2016) | - |

Finally, we analyzed the dominant directional interactions between the seed regions in PFC and the target areas in the visual cortex during the resting state. To this end, we carried out directionality analyses between FEF, IFJa, IFJp, and all the ROIs in Table 1. We computed partial directed coherence (PDC) (Baccalá & Sameshima, 2001) of each unidirectional interaction between seed regions and visual streams. For each pair, we subtracted the PDC values of one unidirectional connection from the other and tested its significance against zero. We chose PDC from a range of directionality measures because it only considers direct interactions between regions, is generally regarded as insensitive to source leakage, and shows high group-level repeatability, for instance, in comparison to the phase slope index (Colclough et al., 2016).

### 2.8 Statistical analyses

We implemented paired Wilcoxon signed-rank tests for each type of analysis because the obtained connectivity values had a non-gaussian distribution. For the same reason, we did not apply z-transformation to the correlation matrix resulting from the oPEC metric. Additionally, we used the false discovery rate (FDR) method to correct for multiple comparisons (Benjamini & Hochberg, 1995; Groppe, 2021). In the hypothesis-driven ROI analyses, we corrected multiple comparisons for all 33 regions in Table 1 (+3 seed regions for PDC measure). Whereas, we constrained the significance test for the whole-brain exploratory analyses to one hemisphere and accordingly corrected multiple comparisons for 180 regions. To visualize FEF and IFJ’s predominant functional connectivities across frequency bands or hemispheres, we converted the adjusted p-values to z-scores. We summed the z-score of each parcel and divided them into the number of times they were significant. Importantly, we applied paired significance tests separately for FEF, aIFJ, and pIFJ per region. We did not average the connectivity values of aIFJ and pIFJ before statistical analyses since we suspected them of having different functional connectivity fingerprints based on previous evidence (Baker et al., 2018; Bedini & Baldauf, 2021).

### 2.9 Ground truth analysis

Lastly, we performed a ground truth analysis to ensure that our analysis pipeline was implemented correctly and provided valid results. We replicated the results from Hipp et al. (2012) paper for the oPEC metric. In the beta band (13-30 Hz), we calculated the power correlation of left and right MT+ with the rest of the brain. We could successfully replicate the seed-based connectivity of MT+ and reproduce the third figure of their paper by using a different dataset, source-reconstruction method (MNE rather than beamforming), and our own processing pipeline. We did not apply any statistical mask to the figure shared in the supplementary material (Figure S1).

## 3 RESULTS

We first report the outcomes for 2 s epoch segmentation and ipsilateral functional connectivity. We detail the results for ROI analyses with respect to phase- and power-based functional connectivity metrics. Additionally, we present PDC in the resting state to expand on the direction of the interaction. The abbreviations for the cortical area names are based on Glasser et al. (2016). The supplementary material provides the findings for 5 and 10 s epoch lengths (Figures S5-6), contralateral connectivity (Figure S9), and whole-brain connectivity analyses (Figures S10-14).

### 3.1 Predominant functional connectivities

All three connectivity metrics demonstrated that FEF has robust functional connectivity with the regions in the dorsal visual stream, whereas IFJa has stronger functional connectivity with the areas in the ventral visual stream independently of lateralization and frequency band (Figure 1; two-sided Wilcoxon signed-rank test, p < 0.05, FDR-corrected). The direct contrast between the ipsilateral connectivity fingerprints of FEF and IFJa (FEF-IFJa) revealed that FEF has a robust power correlation with the dorsal visual stream in beta and gamma bands. In contrast, IFJa has a strong power coupling with the ventral visual stream in delta, beta, and gamma oscillations. Moreover, both iCOH and dwPLI metrics showed that FEF is phase coupled with superior parietal lobule (SPL) and IPS in the beta band, while IFJa is phase-coupled with the inferior and middle temporal cortex by delta and gamma oscillations (Figure 2). Importantly, the anterior and posterior parts of IFJ have different connectivity patterns. Although the direct contrast between the ipsilateral connectivity fingerprints of FEF and IFJp (FEF-IFJp) also indicates FEF’s higher functional connectivity to SPL and IPS, we observed that IFJp has weaker functional connectivity to the middle and ventromedial temporal cortex, particularly in the alpha and beta frequencies. Nevertheless, it still achieves a higher power correlation with the ventral visual areas in the delta frequency compared to FEF (Figures S2 and 2).

**FIGURE 1.**
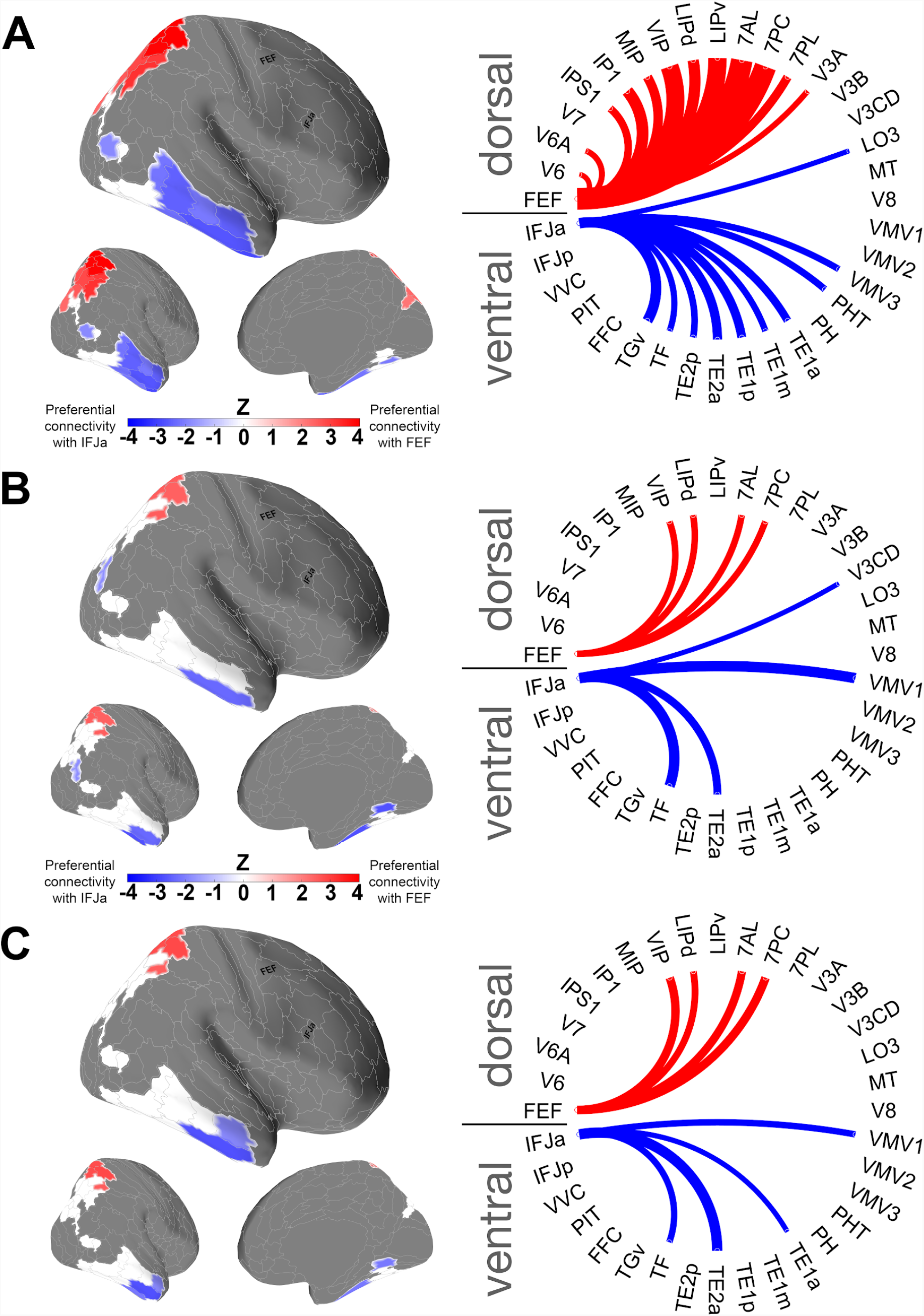
Predominant functional connectivity maps of FEF and IFJa across frequency bands and hemispheres for **(A)** oPEC, **(B)** iCOH, and **(C)** dwPLI metrics (two-sided Wilcoxon signed-rank test, p < 0.05, FDR-corrected for 33 ROIs). The results shown here are based on 2 s epoch segmentation.

**FIGURE 2.**
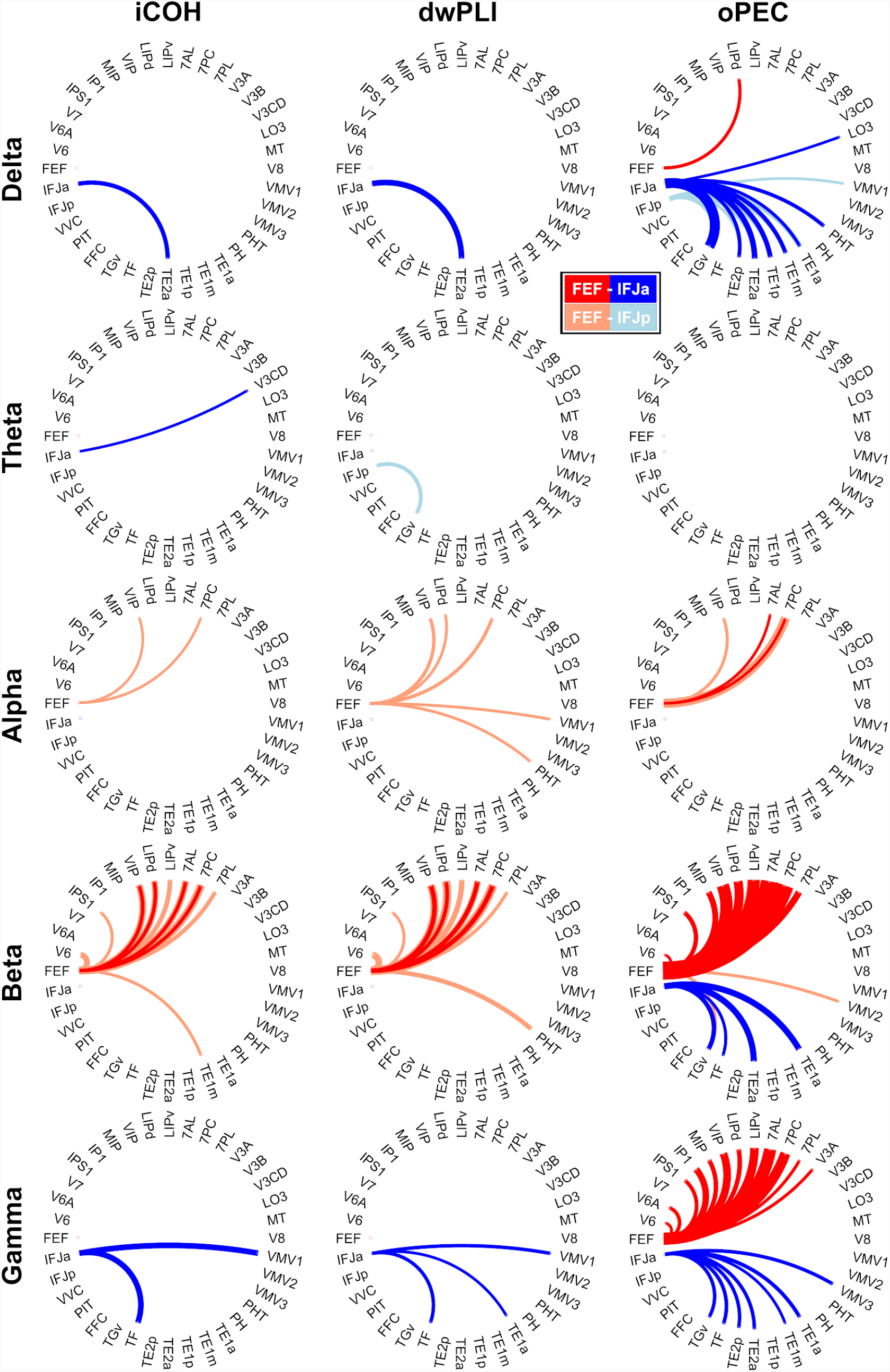
Frequency-specific predominant functional connectivity patterns of FEF, IFJa, and IFJp. The width of a line reflects its relative z-score of the connectivity to other nodes (two-sided Wilcoxon signed-rank test, p < 0.05, FDR-corrected for 33 ROIs). The results shown here are based on 2 s epoch segmentation.

The phase-based metrics showed that the significant parcels for FEF-IFJa are lateralized in the left hemisphere. The iCOH and dwPLI metrics did not reveal any predominant coupling for FEF or IFJa with the selected ROIs in the right hemisphere. On the contrary, while the strong power correlation for FEF-IFJa was evident in both hemispheres, the predominant power coupling between the left IFJa and ventral visual stream was weaker than in the right hemisphere (Figure S3A). Additionally, we observed an apparent dissociation within the higher frequencies for the power coupling between visual streams and plPFC in the right hemisphere. The right FEF and IFJa functionally segregate the right visual pathways into dorsal and ventral streams in beta and gamma oscillations. Overall, the results from both phase- and power-based metrics illustrate that FEF and the areas in the dorsal visual stream predominantly communicate in beta frequency. In contrast, IFJa and the regions in the ventral visual stream are functionally connected through delta and gamma oscillations.

For FEF-IFJp, we observed predominant phase coupling only for FEF in the alpha and beta bands, in the left and right hemispheres, respectively (Figure S3). AdditionallyFurthermore, the iCOH and dwPLI measures showed that areas PHT, TE1m, and VMV1 in the ventral stream have a more robust phase relation with FEF than IFJp. Like the contrast between FEF and IFJa, a predominant power correlation exists in both hemispheres also for FEF-IFJp. However, we cannot see any predominant power coupling between IFJp and the ventral visual stream in higher frequencies. Also, the left VMV2 in the ventral visual pathway has a higher power correlation with the left FEF rather than IFJp.

Noticeably, areas V7, V3B, MT, V8, PH, FFC, PIT, and VVC do not show predominant connectivity either with FEF, IFJa, or IFJp (Figures 1 and S2). In this respect, our additional analyses for FEF, IFJa, and IFJp’s seed-based intrinsic functional connectivities (one-sided Wilcoxon signed-rank test against the average, p < 0.05, FDR-corrected) revealed that V7, V3B, MT, V8, PIT, and VVC do not have statistically significant power or phase coupling with any of these three seed regions (Figure S4). On the other hand, PH has a significant power correlation both with FEF and IFJa. FFC is significantly power coupled with IFJa and IFJp, but not to FEF. Overall, we observed that all three seed regions have a robust power correlation with the parcels in SPL and IPS. They also have statistically significant spontaneous power coupling with the areas in the ventral visual stream.

### 3.2 Influences of chosen epoch length

We observed that 2, 5, and 10 s epoch segmentation affects distinct frequency bands and connectivity metrics differently. In the delta band, IFJa has a consistent robust power correlation with the ventral visual stream. However, its dominance in the ventral pathway decreases in beta and gamma oscillations with increasing segmentation length. In contrast, FEF’s power coupling with the dorsal visual stream is consistent in beta and gamma frequencies (Figure S5). Additionally, the results for iCOH are less consistent as the epoch length increases and the number of trials decreases per recording session (Figure S6). Nevertheless, specific predominant functional couplings stand out across different epoch segmentation and connectivity measures: FEF has robust functional connectivity with the areas VIP, LIPd, 7AL, and 7PC in the dorsal visual stream in higher frequencies. Whereas IFJa is predominantly coupled with the areas TE1, TE2, TF, and TGv in the ventral visual stream in lower frequencies.

### 3.3 Directional interactions between visual streams and plPFC

Directionality analyses in the frequency domain revealed intrinsic unidirectional information flows between seed regions and the visual streams (Figure 3; two-sided Wilcoxon signed-rank test, p < 0.001, FDR-corrected). Most remarkably, PDC showed that directional interactions from SPL and IPS to seed regions are significantly more potent than in the other direction. In contrast, directional influences from seed regions to the middle and inferior temporal cortex are more robust than in the other direction, mainly in delta, theta, and beta frequency bands. In other words, the rMEG time series of the regions in plPFC is predicted by the intrinsic neural activity of the regions in the superior parietal cortex (SPC; SPL and IPS) and themselves predictive of the spontaneous activity of the areas in the ventral visual stream. In the alpha band, we instead observed that the seed regions, particularly IFJa and IFJp, predict the intrinsic neural activity in both visual streams except areas 7AL, 7PL, and VIP in the dorsal stream. The direction of information flow from seed regions to these areas located in the medial part of SPC (mSPC) is not significantly stronger than the other direction for any oscillatory band. Likewise, no region in the ventral stream has a directional influence over any seed region, except left TE1m in the gamma band.

**FIGURE 3.**
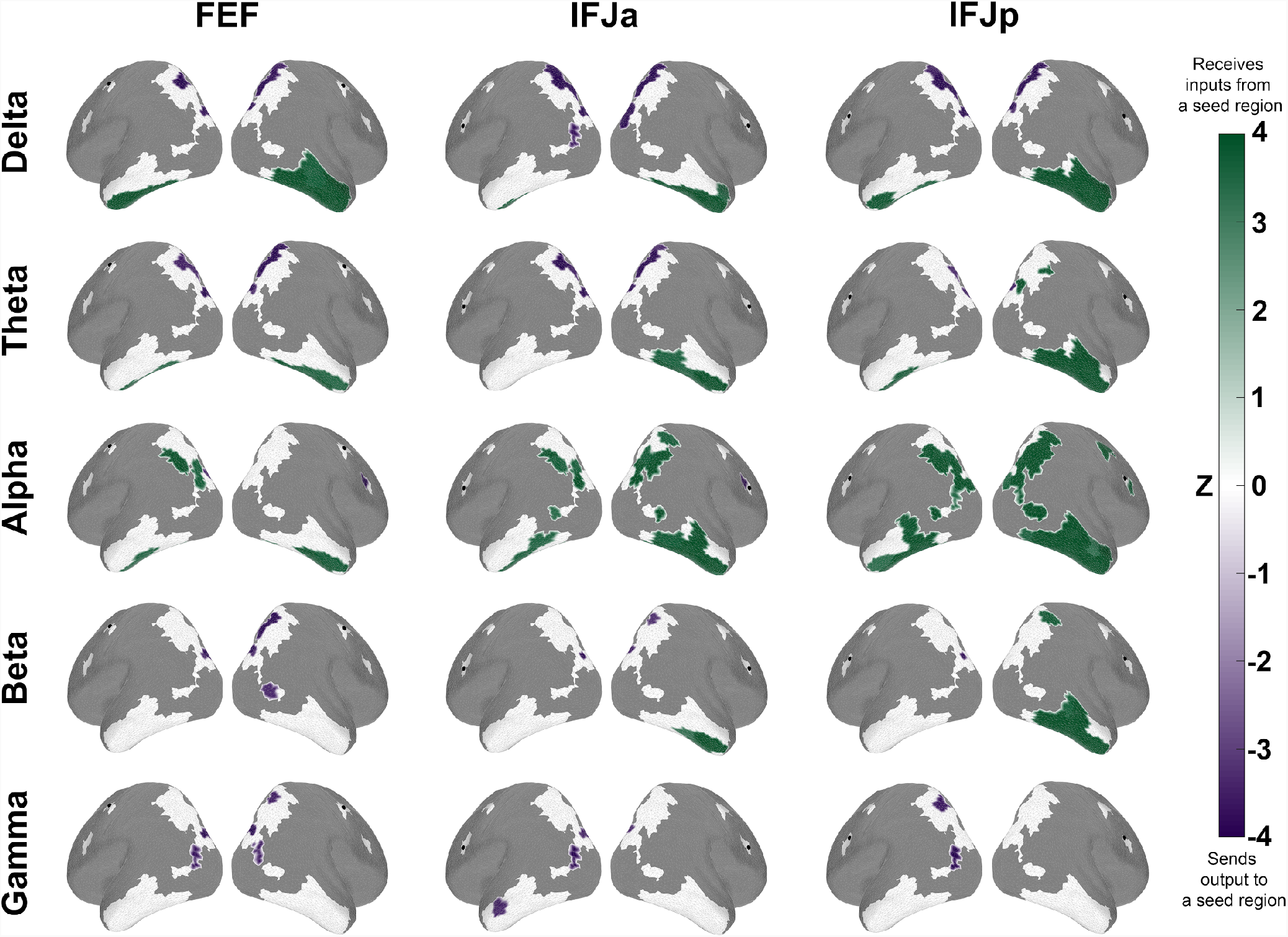
Partial directed coherence between FEF, IFJ, and visual streams. The figure shows the directional influences between seed regions (FEF, IFJa, and FEF) and visual streams. While the green regions receive inputs, the magenta ones send outputs to a seed region (two-sided Wilcoxon signed-rank test, p < 0.001, FDR-corrected for 33+3 ROIs). The results shown here are based on 2 s epoch segmentation.

### 3.4 Directional Interactions Among FEF, IFJa, and IFJp

Finally, the directionality analyses between FEF, IFJa, and IFJp indicated a direction of information flow from IFJp to the other two seed regions (Figure 4). The PDC values from IFJp to IFJa and IFJp to FEF are statistically stronger than in the other directions in both hemispheres. While left IFJp feeds information to left FEF and left IFJa mainly in the theta band, the same direction of interaction in the right hemisphere is driven through both alpha and beta oscillations. Notably, the directed interaction from right IFJp to right FEF in the alpha band is highly significant (p < 0.00001; Figure S7). Besides, we observe that right FEF and IFJa have reciprocal connections in various frequency bands. The direction of spontaneous activity from FEF to IFJa and IFJa to FEF are subserved by delta and alpha oscillations, respectively. In the left hemisphere, we only see unidirectional interactions from FEF to IFJa in the delta and gamma band but not vice versa. Noticeably, the delta frequency band underlies FEF’s directional influence over IFJa in both hemispheres (p < 0.005).

**FIGURE 4.**
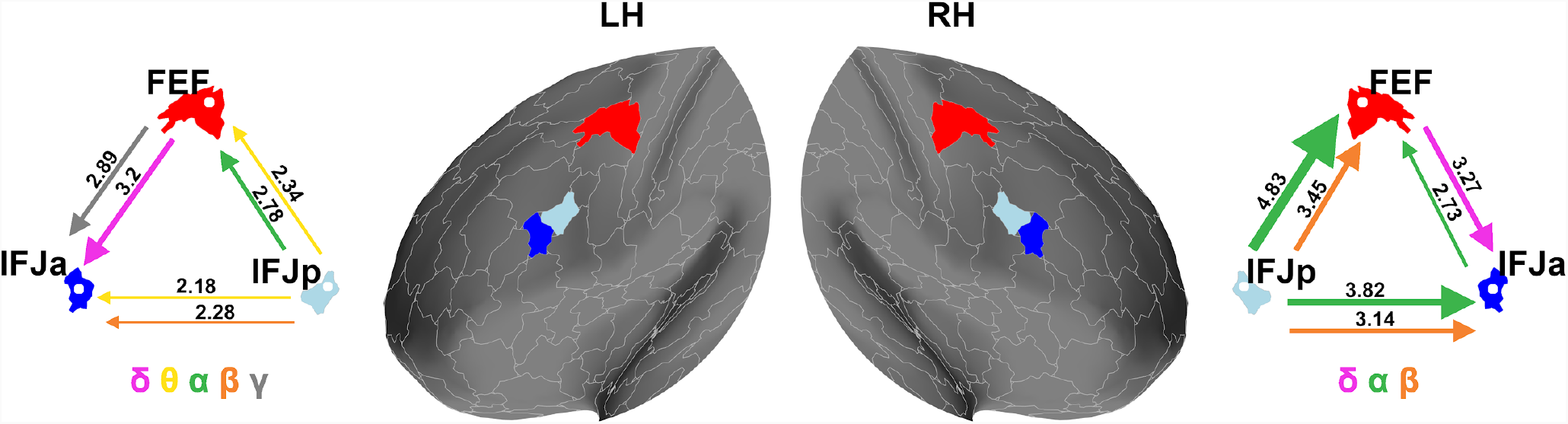
Partial directed coherence between the seed regions within PFC. The figure shows the directional influences among FEF, IFJa, and IFJp in both hemispheres. The width of the arrows reflects z-scores on top of them (two-sided Wilcoxon signed-rank test, FDR-corrected for 33+3 ROIs; see Figure 4). The distribution of subjects’ PDC value for each significant directional influence is illustrated in Figures S7 and S8.

## 4 DISCUSSION

The current study investigated the intrinsic functional connectivity profiles of FEF and IFJ, two specific control circuits in the dorsal and ventral PFC, to provide additional evidence for their respective top-down functions in spatial versus non-spatial (feature and object-based) attention and working memory control. Our results demonstrated that FEF has a more robust power and phase coupling with the spatiotopically organized regions in the dorsal visual stream. In contrast, IFJa has more substantial power and phase relation with the feature or object encoding regions in the ventral visual pathway (Figure 1). These results suggest that in humans the ‘where’ and ‘what’ visual streams (Goodale & Milner, 1992; Mishkin et al., 1983) might extend all the way into plPFC (Goldman-Rakic, 1996; Macko et al., 1982; Wilson et al., 1993). The spatial versus non-spatial to-be-attended or to-be-remembered stimulus domain might respectively segregate plPFC into a functionally specific dorsal versus ventral visual pathways.

### 4.1 FEF vs. IFJa

FEF has robust functional connectivity with the areas VIP, LIPd, 7AL, and 7PC across different epoch lengths and connectivity metrics in the dorsal stream. In this respect, studies on monkeys reported that area VIP is involved in the analysis of visual motion (Colby et al., 1993; Duhamel et al., 1998), multisensory space representation (Schlack et al., 2005), visuomotor control (Culham et al., 2006), self-motion (Bremmer, 2005; Chen et al., 2011; Schlack et al., 2002) and motion encoding in near-extrapersonal space (Schlack et al., 2003). While area LIP is also to some degree selective to shape (Sereno & Maunsell, 1998), researchers at large showed its role in spatial attention (Bisley & Goldberg, 2003; Thakral & Slotnick, 2009), the choice of saccade target (Andersen et al., 1990; Barash et al., 1991; Blatt et al., 1990; Kusunoki et al., 2000; Kusunoki & Goldberg, 2003) and the creation of priority maps (Gottlieb, 2007; Ipata et al., 2009; Sprague & Serences, 2013). Finally, it is evident that SPL (areas 7AL and 7PC) takes part in shifting spatial attention (Behrmann et al., 2004; Caspari et al., 2018; Corbetta et al., 1995; Kelley et al., 2008; Molenberghs et al., 2007; Spadone et al., 2021; Vandenberghe et al., 2001; Yantis et al., 2002) and encoding spatial coordinates for planned reach movements (Baldauf et al., 2008; Connolly et al., 2003). Besides, these results match with FEF’s deterministic tractography (Baker et al., 2018). FEF’s white matter connections accordingly terminate at the intraparietal sulcus (IP1 and IP2), nearby SPL. Overall, the spontaneous connectivity between FEF and SPC (SPL and IPS) corresponds to their underlying structural connectivity and functional specialization in spatial attention and working memory.

In contrast to FEF, IFJa has a stronger functional coupling with the areas TE1a, TE2a, TF, and TGv in the ventral visual stream. These areas roughly correspond to the anterior middle temporal gyrus (aMTG; area TE1a), inferotemporal cortex (IT; areas TE2a and TF) and temporal pole (TP; area TGv). Notably, studies associated aMTG with social cognition (Xu et al., 2020), language, and semantic processing (Ferreira et al., 2015; Gorno-Tempini & Price, 2001; Rossell et al., 2001; Siuda-Krzywicka et al., 2021; Walker et al., 2011). On this note, this particular functional coupling with the anterior temporal cortex might be due to mind-wandering during the resting-state scan (Chou et al., 2017). In comparison, lesions and neuroimaging studies suggest that TP takes part in face recognition by associating person-specific stored memories to the perceptual representation of people’s faces (DeVries & Baldauf, 2019; Kamps et al., 2019; Landi et al., 2021; Olson et al., 2007). Additionally, a relational processing task adapted from Smith et al. (2007) revealed that, in contrast to TGd, TGv is activated vs deactivated in relational primary contrasts when subjects were asked to differentiate objects based on their shapes or textures (Glasser et al., 2016). Lastly, the IT cortex has a direct role in object recognition (Kreiman et al., 2006; Lehky & Tanaka, 2016; Tanaka, 1996; Yamane et al., 2008). Studies on monkeys and humans both show IT cortex’s involvement in object-based attention and working memory (Baldauf & Desimone, 2014; Kar et al., 2019; Miller et al., 1996). Furthermore, IFJa, but not FEF, has distinctive white matter connectivity with the temporal cortex, which is in accordance with its predominant functional coupling reported here. IFJa’s deterministic tractography demonstrates its structural connectivity to the middle and temporal gyrus (Baker et al., 2018). Also, IFJ’s probabilistic DWI tractography is especially well connected with several functional areas in IT cortex, which was not the case for FEF (Baldauf & Desimone, 2014). All in all, in light of previous evidence, we indicate that FEF and IFJa’s respective predominant functional coupling with SPC and the temporal cortex in the resting state is congruent with their respective functional segregations in plPFC.

### 4.2 Frequency-specific interactions

Different oscillations are associated with different cognitive functions (Başar et al., 2001; Güntekin & Başar, 2016; Harmony et al., 1996). Thanks to the high temporal and spatial resolution of MEG, we could reveal the frequency-specific signatures of the functional coupling between FEF, IFJ, and visual pathways (Figure 2). While these frequency-specific interactions in resting state data hint at general communication patterns between the involved areas, it is important to bear in mind that those interactions can rapidly change either in strength, dominant frequency band, and/or directionality as specific attentional or working memory tasks evolve.

In rest, we found substantial delta oscillatory coupling between IFJ and the areas in the ventral visual stream. In this respect, Harmony and colleagues (1996) observed that attention to internal representations leads to increased delta power during mental processing, such as the Sternberg paradigm that measures working memory performance. They suggested that delta oscillations carry attentional modulations from the frontal cortex to distant neuronal networks (Harmony, 2013). Such that it might have a role in suppressing internal or external distractors that interfere with the ongoing mental state. Noticeably, these statements align with IFJ’s role in cognitive control (Brass et al., 2005; Derrfuss et al., 2005) and selective attention (Baldauf & Desimone, 2014; Bedini & Baldauf, 2021). Also, for example, Zarjam et al. (2011) could successfully classify working memory load levels based on the delta activity in the frontal EEG. Moreover, they linked delta synchronization to subjects’ internal concentration. Since functional coupling between distant regions requires a slower ‘clock’ (Jensen & Colgin, 2007), delta oscillations might subserve the intrinsic functional connectivity between IFJ and the areas in the ventral visual stream in the absence of attention to features or objects. This baseline information flow can potentially pave the way for task-evoked activity (Cole et al., 2016) during selective attention and working memory.

In the theta band, we did not find any significant region that predominantly power correlates with FEF or IFJ. Additionally, phase-based measures did not reveal any consistent results. The lack of predominant coupling in the theta band might be because theta oscillations are instead linked to learning and memory through cortico-hippocampal interactions (Başar et al., 2001; Buzsáki, 2002; Goutagny et al., 2009; Herweg et al., 2020; Lega et al., 2012; Lubenov & Siapas, 2009; O’Neill et al., 2013; Siapas et al., 2005). However it is important to note that several task-based studies linked theta oscillation to visual working memory (Liebe et al., 2012; Muthukrishnan et al., 2020; Sauseng et al., 2010) and willed attention (Rajan et al., 2019).

On the other side, we found predominant functional connectivity between FEF and SPC (areas VIP, LIPd, 7AL, and 7PC) in the alpha band. Some of this alpha band activity in the dorsal connections, might be attributed to general arousal (Barry et al., 2020; Brancaccio et al., 2020; Cantero et al., 1999; Pivik & Harman, 1995) or visual mental imagery (Xie et al., 2020) during resting state. However, alpha oscillations are also known as neural signature in task-based attention studies (Babiloni et al., 2006; Bae & Luck, 2019; Bagherzadeh et al., 2020; Capotosto et al., 2009; DeVries et al., 2021, 2019; Geng, 2014; Klimesch et al., 2007; Mazaheri et al., 2011; Noah et al., 2020; Rihs et al., 2007; Sauseng et al., 2005; Thut et al., 2006; Toscani et al., 2010; VanDiepen et al., 2016).

Beta synchronization, which we found more strongly involved in the dorsal connections of FEF, is associated with motor inhibition (Gilbertson et al., 2005; Picazio et al., 2014) and termination (Heinrichs-Graham et al., 2017; Kilavik et al., 2013; Pfurtscheller et al., 1996). In contrast, its desynchronization is linked to movement preparation and execution (Kühn et al., 2004; Pfurtscheller et al., 1997; Pfurtscheller & Lopes da Silva, 1999; Y. Zhang et al., 2008). Even so, the entrainment of beta-band activity through transcranial alternating-current stimulation could slow down voluntary movements in healthy subjects (Pogosyan et al., 2009). In this respect, Engel and Fries (2010) suggested that beta oscillations signal the status quo, i.e., the maintenance of the current sensorimotor set. They claimed that the intention or prediction to hold on to the present state leads to robust beta band coupling. Accordingly, we reason that the robust beta oscillatory coupling between FEF and SPC stems in part from the suppression of eye movements during fixation on the crosshair. It might reflect the inhibition of the oculomotion, sustained attention, and the status quo during resting state. Thereby, movement or visuomovement cells in FEF (Gregoriou et al., 2012) likely drive FEF’s intrinsic beta synchronization with SPC. Moreover, the deficits in motor response inhibition are linked to attention deficit hyperactivity disorder (ADHD) (Mostofsky et al., 2001). Compared to control subjects, people with ADHD perform particularly poorly in anti-saccade tasks that require suppressing reflexive eye movements. They also have difficulties maintaining steady fixations for a prolonged time (Munoz et al., 2003; Munoz & Everling, 2004). Consequently, it has been argued that the disruption in FEF and SPC’s functional coupling in the beta band might indicate a specific deficit in endogenous visuospatial attention (Caldani et al., 2020).

Engel and Fries (2010) proposed that while beta-band activity signals the maintenance of the status quo, gamma-band oscillations indicate the readiness to change it. In line with this statement, several studies reported task-induced gamma synchrony in object perception, attention, and working memory (Baldauf & Desimone, 2014; Fries, 2005, 2009, 2015; Gregoriou et al., 2009; Howard et al., 2003; Tallon-Baudry et al., 1998; Tallon-Baudry & Bertrand, 1999). Despite its specific role in the task-based activity, all three connectivity measures of our present analysis revealed spontaneous functional connectivity between IFJ and the areas in the ventral visual stream in the gamma band. We suggest that, in particular, IFJ and IT cortex’s intrinsic gamma coupling hints at their task-evoked functional connectivity during object-based attention (Baldauf & Desimone, 2014). However, it is also likely that this spontaneous synchronization between IFJ and the ventral stream is driven by mind-wandering or imagery in the scanner (Albers et al., 2013; de Borst et al., 2012; Dentico et al., 2014; Dijkstra et al., 2017, 2018; Pearson, 2019). Finally, the predominant functional coupling between IFJ and the ventral visual stream in the rest might to some extent also be driven by cross-frequency coupling of delta and gamma rhythms (Bruns & Eckhorn, 2004; Canolty & Knight, 2010).

### 4.3 Directionality of interactions

In addition to bidirectional functional coupling, we investigated unidirectional interactions between plPFC and visual streams in the resting state. The majority of our regions of interest in Table 1 exhibited a clear directionality to the respectively connected seed region in the lower frequencies. Remarkably, partial directed coherence revealed directed interactions from mSPC (areas 7AL, 7PL, and VIP) to FEF and IFJ in delta and theta oscillations (Figure 3). Since in the delta and theta range the neural time series of mSPC are statistically more successful in predicting the neural activity in FEF and IFJ than vice versa, we can suggest that SPC drives the spontaneous activity in plPFC during rest. This unidirectional connectivity during rest might reflect similar task-related influences of SPC over plPFC during attention and working memory (Chen et al., 2020; Spadone et al., 2021). Additionally, we observe a frequency-specific direction of interaction in the beta band from parietal areas VIP and 7PL to FEF. A recent study by Spadone et al. (2021) similarly indicated that contrasting stay with shift cues reveals a robust direction of information flow from the right ventral IPS to the right dorsal FEF, specifically in the beta band. Since subjects had to fixate during rMEG, the maintenance of attention and the status quo of beta oscillations could have led to the results we observed in the current study. The feedback influences of IPS onto FEF in the beta band might signal to sustain eye movements on the crosshair. Finding such directionality from the parietal cortex to FEF in the beta band is somewhat surprising and speaks against the general notion that the beta band solely provides top-down signals. For these reasons, it is important to note that the direction of an oscillatory coupling is often task-dependent. Therefore, the direction of interaction between SPC and FEF might change depending on the current task demands. While SPC’s highly significant directional influence over plPFC during rest suggests that it might provide some sort of spatial attention signals to FEF, Spadone et al. (2021) also showed that the contrast between shift versus stay cues reveals feedback connections from the right FEF to the right SPL. In this respect, the functional hierarchy is not fixed but dynamic. Additionally, the directionality measures should be taken cautiously in general since unidirectional interactions can be observed also because of the asymmetries in the noise levels (Nolte et al., 2004).

### 4.4 IFJa vs. IFJp

The functional connectivity analyses (FEF-IFJa and FEF-IFJp) did not reveal identical predominant intrinsic connectivities for IFJa and IFJp (Figures 1 and S2). In particular, IFJa has generally a stronger functional coupling with the ventral visual stream than IFJp. Since these areas have different intrinsic connectivity fingerprints, we suggest that they are functionally distinct regions (Passingham et al., 2002), as already emphasized in MMP1 (Glasser et al., 2016) and previous studies. For instance, Zanto et al. (2011) pointed out that the right IFJ’s dorsal and ventral parts are respectively involved in motion and color processing (see also Schwedhelm et al., 2020). Moreover, Asplund et al. (2010) proposed that IFJ might have a role in both stimulus-driven and goal-directed attention through its participation in the ventral and dorsal attention networks. Furthermore, the deterministic tractography revealed distinct white matter tracts for IFJa and IFJp. IFJa had more robust structural connectivity with the middle and inferior temporal gyrus (areas TE1a, TE1m, and TE2a) compared to IFJp (Baker et al., 2018). Finally, the distinct predominant connectivities of IFJa and IFJp provides strong evidence against the notion that our connectivity results could potentially suffer from a proximity bias (since IFJa is more distant from the visual cortex than IFJp).

We found a highly significant direction of interaction from IFJp to FEF and IFJa in both hemispheres (Figure 4). Within the right hemisphere, alpha and beta oscillations govern these intrinsic directional influences from IFJp to the other seed regions. In the left hemisphere, the same direction of information flow is supported also by theta, in addition to alpha and beta bands. Although the direction of interaction may change depending on task-related activity, it is clear that IFJp feeds information to FEF and IFJa in the resting state. Taken together with the previously mentioned result that IFJp has overall less functional connectivity with the ventral visual cortex, this may hint at its role in providing precursor activity to the output units in FEF and IFJa, which then channel that information to visual cortex. Interestingly, also in non-human primate recordings similar interactions have been observed between VPA (the non-human homologue of IFJ) and FEF in visual search tasks: signals from VPA to FEF helped guiding eye movements to likely targets, and the inactivation of VPA selectively eliminated feature attention signals in FEF (while leaving spatial attention and eye movement control unaffected), such that the animals could no longer find targets during visual search (Bichot et al., 2019).

### 4.5 Power- and phase-based connectivity

Power- and phase-coupling potentially reflect partly distinct neuronal mechanisms (Daffertshofer et al., 2018; Siems & Siegel, 2020). While power-based metrics compute functional connectivity between two neuronal oscillations based on their amplitudes’ envelope correlation, phased-based measures instead consider the consistent relative phase difference between them (Siegel et al., 2012). In this sense, phase-based measures such as iCOH or dwPLI are more delicate and more conservative while attributing a genuine functional coupling in comparison to power-based metrics to avoid Type I errors (Nolte et al., 2004; Stam et al., 2007; Vinck et al., 2011). However, they both have also been found to be less replicable on the group level and less consistent within and between subjects in resting state data (Colclough et al., 2016). On the other hand, the orthogonalized power envelope correlations (Brookes et al., 2011; Hipp et al., 2012) are shown to be the most replicable methods among several alternative metrics, and, importantly, they are insensitive to spurious coupling stemming from field-spread (Colclough et al., 2016; Duan et al., 2021). In light of these findings, we state that the robustness of power- and phase-based measures in the current study relates to the nature of these metrics’ computation. Compared to power correlation, the less number of significant phase-couplings for iCOH and dwPLI might be due to their detailed measurement. Meanwhile, the robust connectivity results for oPEC seem to be more reproducible and generalizable across the population. Our successful replication of the results in Hipp et al. (2012) is an additional proof in this respect.

## 4 CONCLUSION

We found that the areas in the dorsal visual stream have a more robust functional coupling with FEF, particularly in the beta oscillations. In contrast, the regions in the ventral visual stream have statistically greater functional connectivity with IFJa, particularly in delta and gamma bands. In light of these and previous findings (Cohen et al., 2008; Cole et al., 2014; Fox et al., 2006; Glasser et al., 2016; Mars et al., 2011; Power et al., 2014; Smith et al., 2009; Tavor et al., 2016; Yeo et al., 2011), we argue that intrinsic connectivity patterns are congruent with each brain region’s function. Therefore, we conclude that FEF’s functional connectivity with the spatiotopically organized regions in SPL and IPS suggests its role in spatial attention and working memory. In contrast, IFJa’s functional coupling with feature- or object-encoding areas in the temporal cortex indicates its role in non-spatial attention and working memory. Our functional connectivity data therefore provides strong evidence that also the prefrontal cortex is organized in a dorsal and ventral stream (Goldman-Rakic, 1996).

The resting-state neuroimaging data is time- and cost-efficient since it eases the acquisition of large datasets and enables analyses of multiple hypotheses over task-free recordings. Nevertheless, it is vital to support the findings from resting-state recordings with task-based settings. To this end, the current study attempted to provide the first direct contrast between the functional connectivity profiles of FEF and IFJ to prompt further research on these regions’ functional integration with the rest of the brain. Future studies should directly compare the functional connectivity fingerprints of these regions in task designs as in Meyyappan et al. (2021) to provide a more complete picture. At the same time, it is also crucial to reveal their underlying structural connectivity patterns, ideally by using probabilistic rather than deterministic tractography since the latter underestimates the existing long-range white matter tracts. Overall, the direct contrast between the intrinsic, functional, and anatomical ‘connectional fingerprints’ (Passingham et al., 2002) of FEF and IFJ would provide further evidence for their functional integration with other cortical regions and a dorsal versus ventral functional segregation within plPFC as an extension of ‘where’ and ‘what’ visual pathways.

## ACKNOWLEDGMENTS

This research is supported by Ministero degli Affari Esteri e della Cooperazione Internazionale (to O.S.) and Fondazione Cassa Di Risparmio Di Trento E Rovereto (to D.B).

## AUTHOR CONTRIBUTIONS

OS preprocessed and analyzed MEG recordings, wrote the initial draft for the manuscript, and prepared the table and figures. DB conceived the study, supervised the data analyses, writing process and the preparation of the figures, and revised the final manuscript.

## DATA AVAILABILITY STATEMENT

The data used in the current study provided and publicly made available by the 1200 Subjects Release of Human Connectome Project (Larson-Prior et al., 2013; Van Essen et al., 2013) in ConnectomeDB (db.humanconnectome.org; Hodge et al., 2016).

