## Supplementary Materials for "Functional connectivity fingerprints of the frontal eye fields and inferior frontal junction in the dorsal vs. ventral prefrontal cortex"

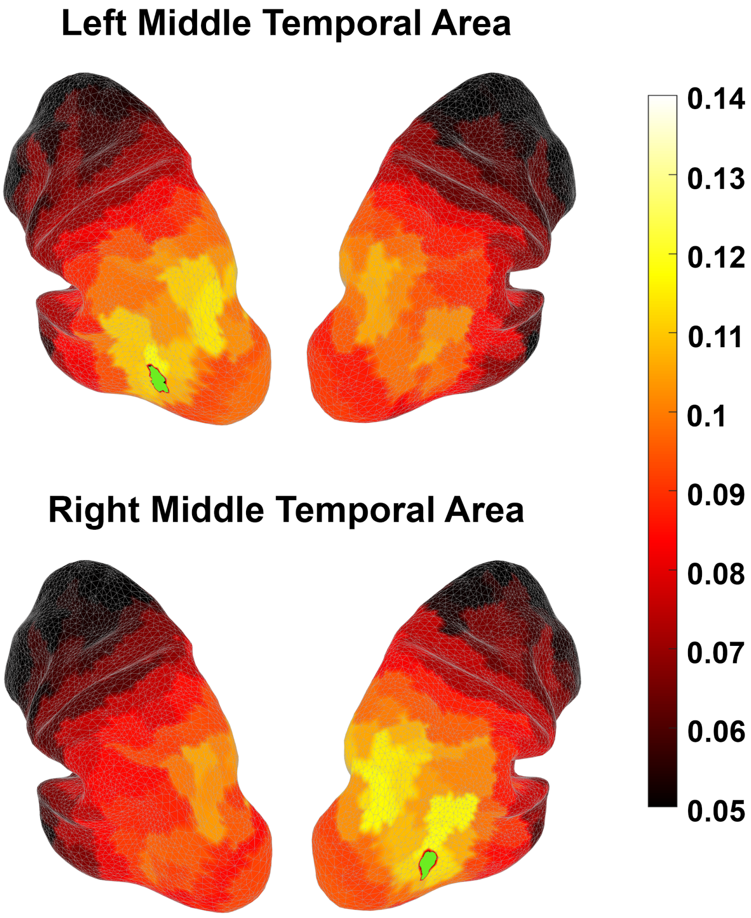


FIGURE S1 The ground truth analysis for the orthogonalized Power Envelope Correlation (oPEC) metric. The figure shows the seed-based intrinsic functional connectivity patterns of the left and right middle temporal area (MT+) in the beta band, replicating figures 3a and 3b in Hipp et al. (2012) paper. We did not apply any statistical masks. The green area illustrates the seed region.

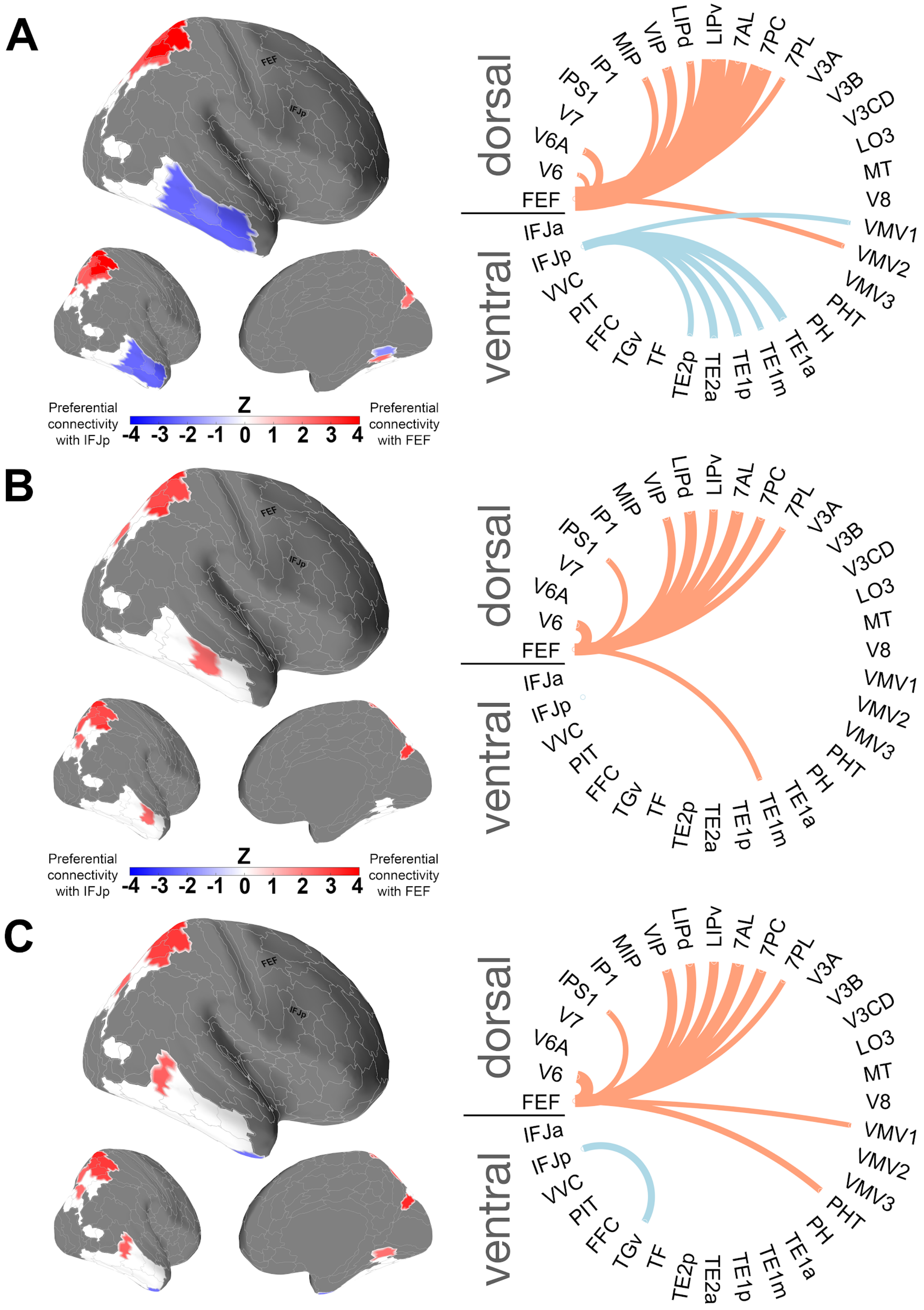


FIGURE S2 Predominant functional connectivity maps of FEF and IFJp across frequency bands and hemispheres for (A) oPEC, (B) iCOH, and (C) dwPLI metrics (two-sided Wilcoxon signed-rank test, $p < 0.05$, FDR-corrected for 33 ROIs). The results shown here are based on 2 s epoch segmentation.

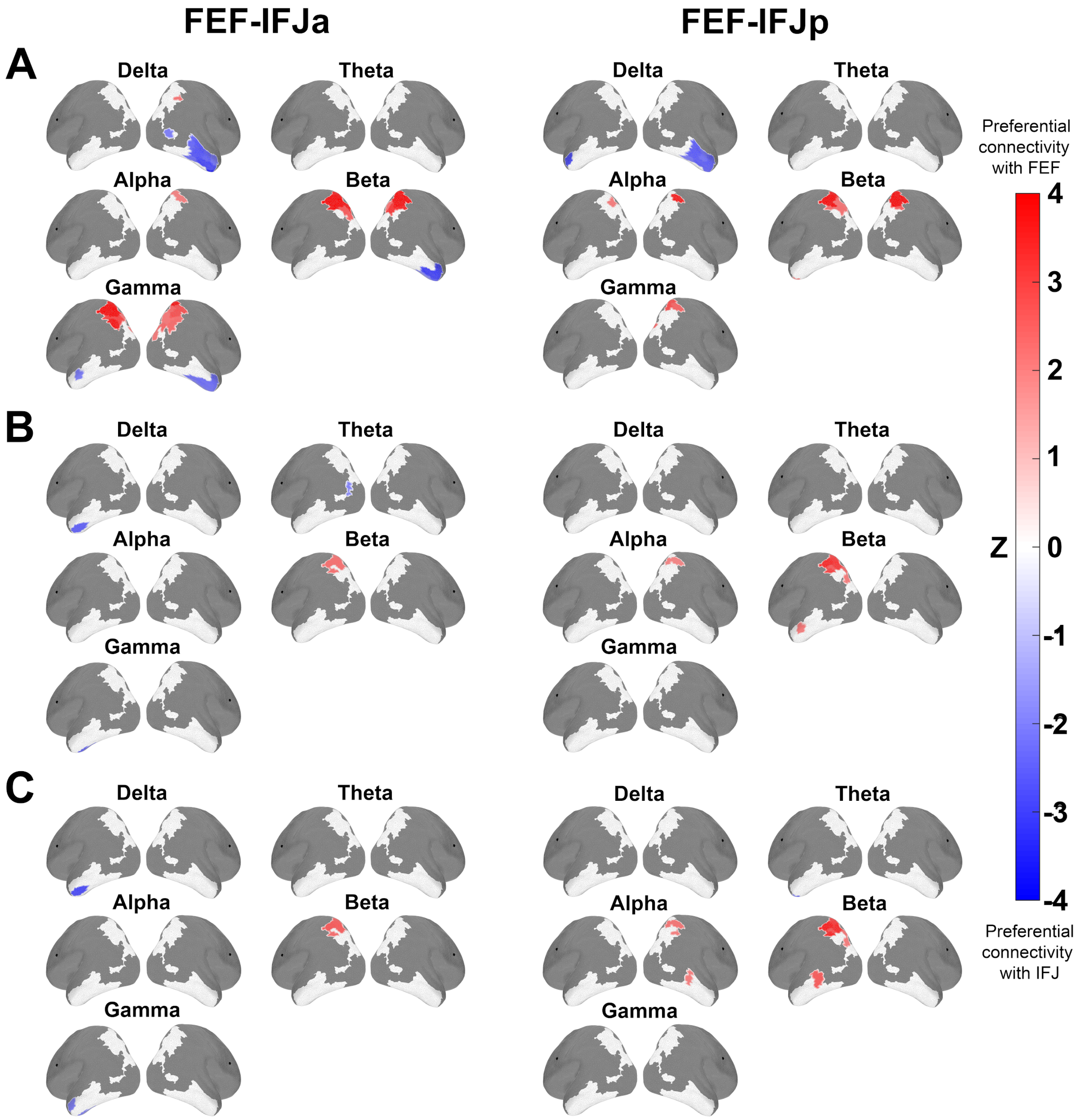


FIGURE S3 The predominant functional connectivity maps of FEF, IFJa, and IFJp per hemisphere for (A) oPEC, (B) iCOH, and (C) dwPLI metrics (two-sided Wilcoxon signed-rank test, $p < 0.05$, FDR-corrected for 33 ROIs). The results shown here are based on 2 s epoch segmentation.

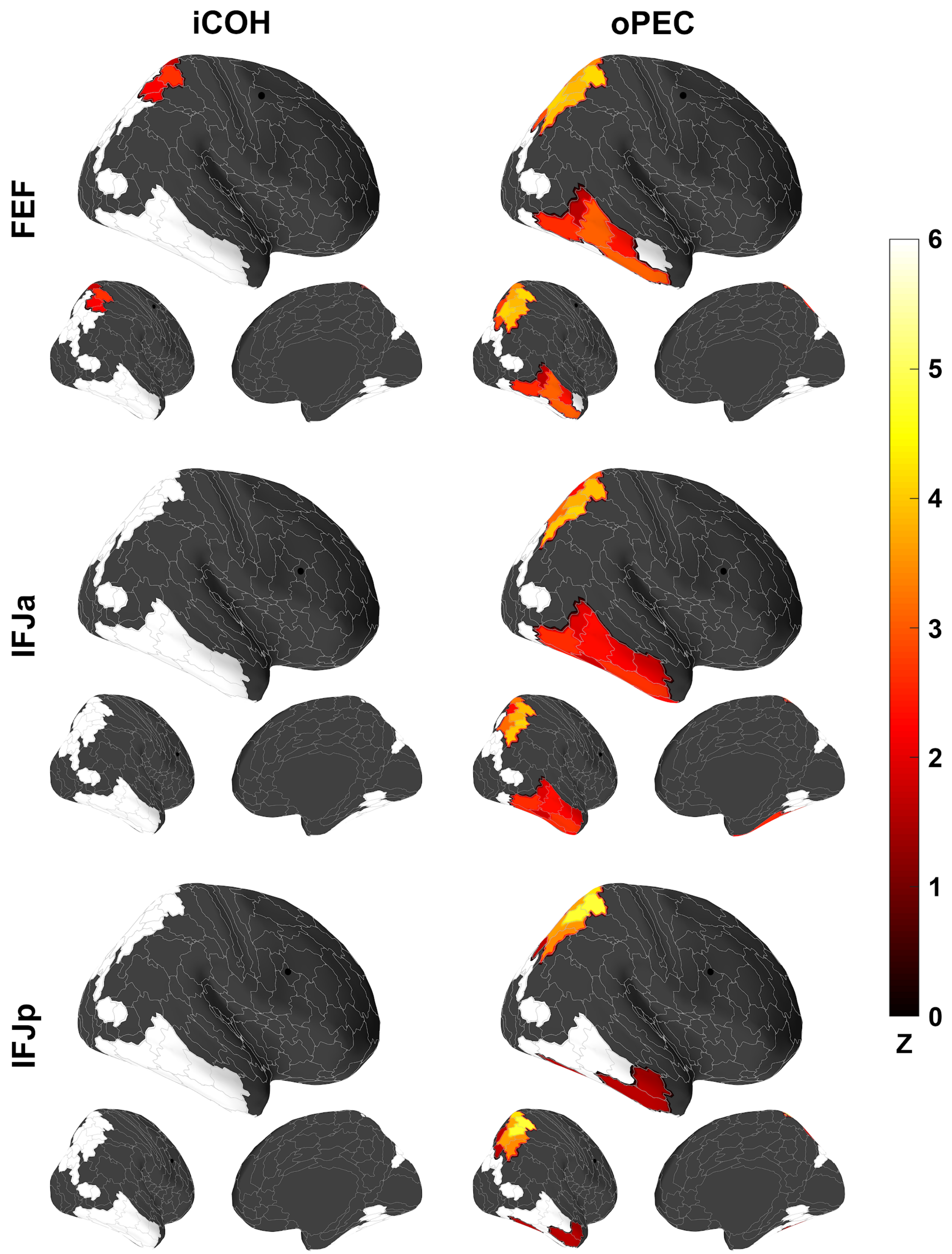


FIGURE S4 Intrinsic functional connectivity profiles of FEF, IFJa, and IFJp for oPEC and iCOH metrics (one-sided Wilcoxon signed-rank test, $p < 0.05$, FDR-corrected for 33 ROIs). The results shown here are based on 2 s epoch segmentation.

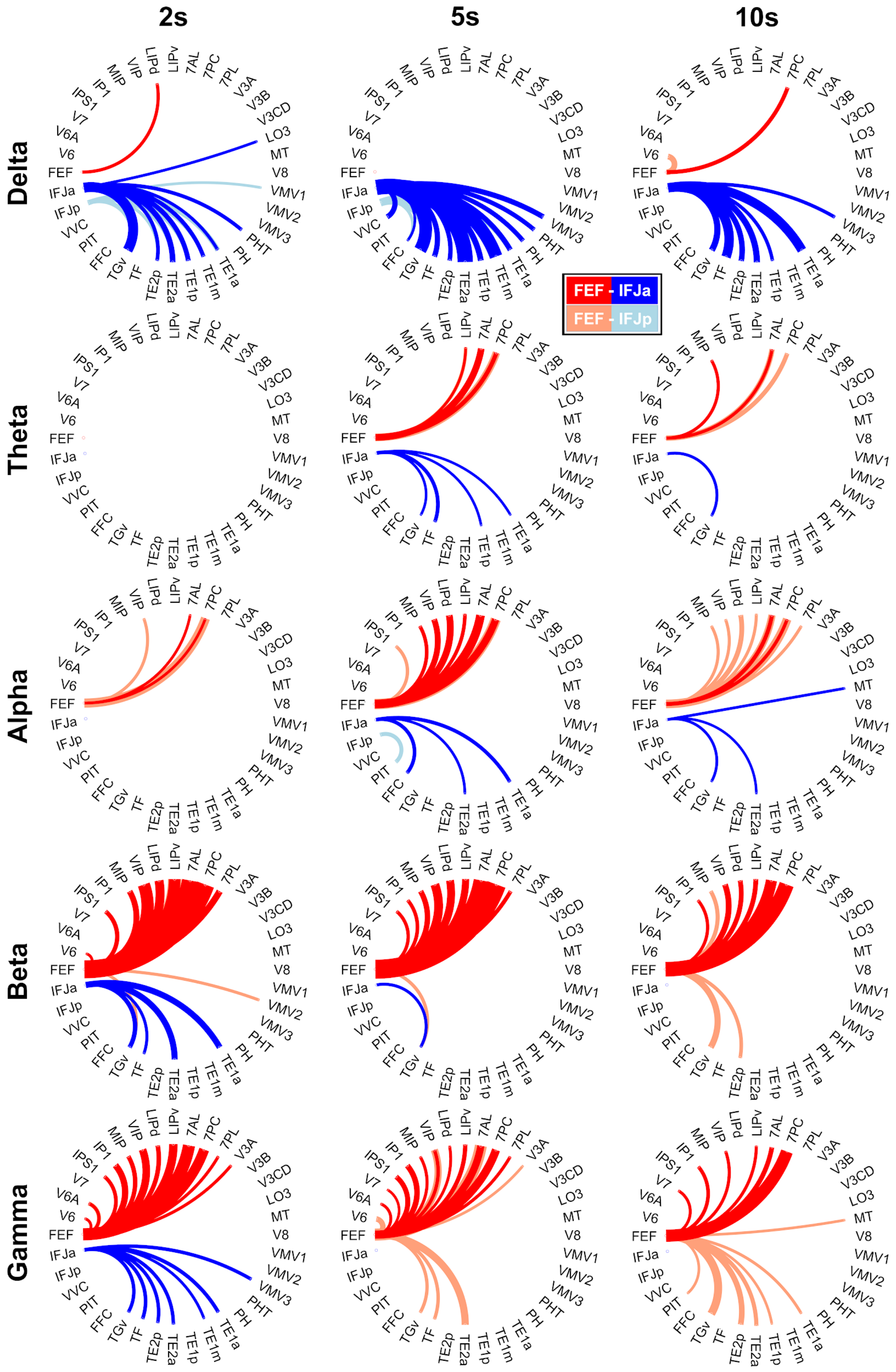


FIGURE S5 The epoch length's impact on the predominant power correlation (oPEC metric) of FEF, IFJa, and IFJp (two-sided Wilcoxon signed-rank test, $p < 0.05$, FDR-corrected for 33 ROIs).

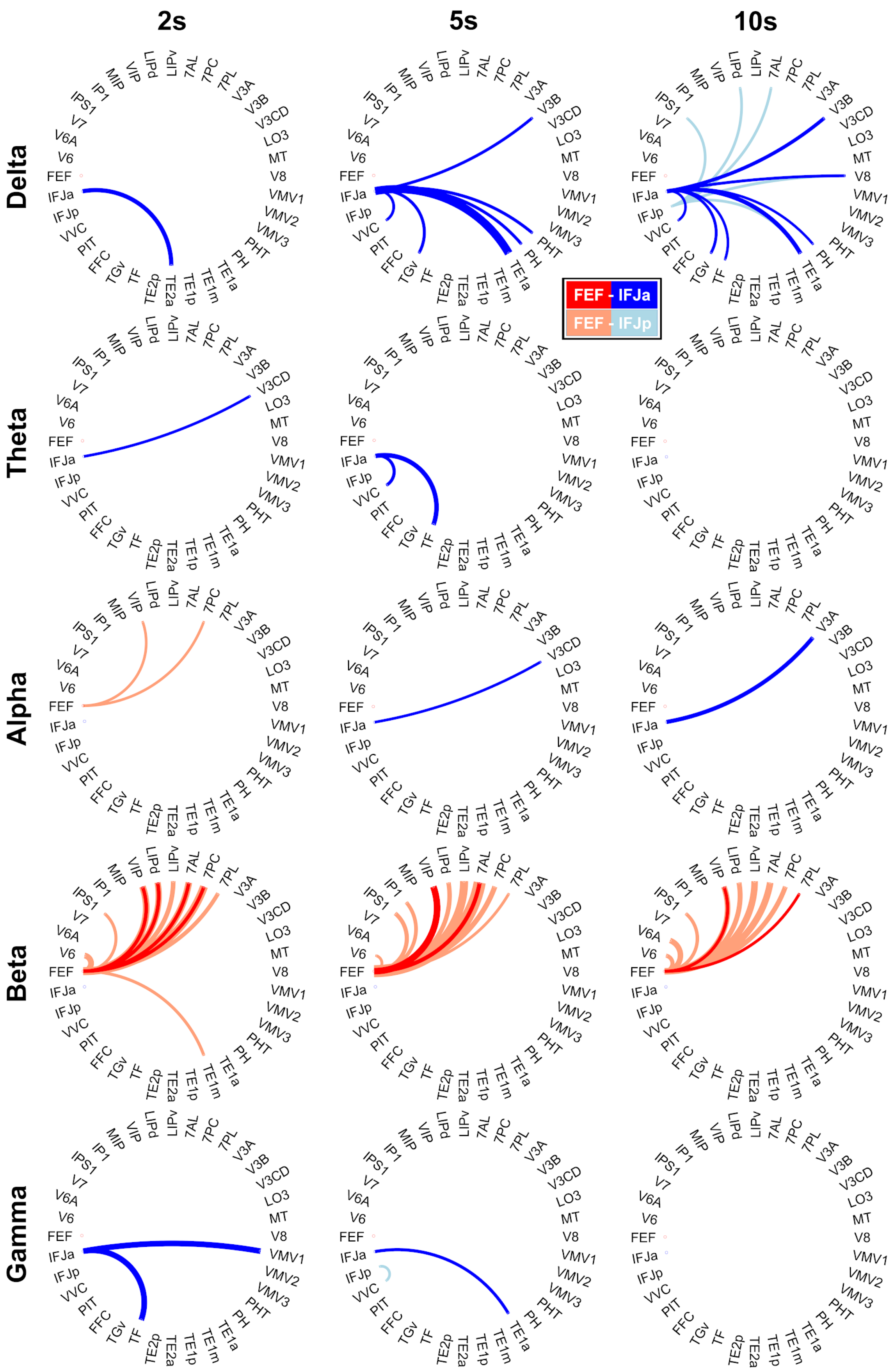


FIGURE S6 The epoch length's impact on the predominant phase coupling (iCOH metric) of FEF, IFJa, and IFJp (two-sided Wilcoxon signed-rank test, $p < 0.05$, FDR-corrected for 33 ROIs).

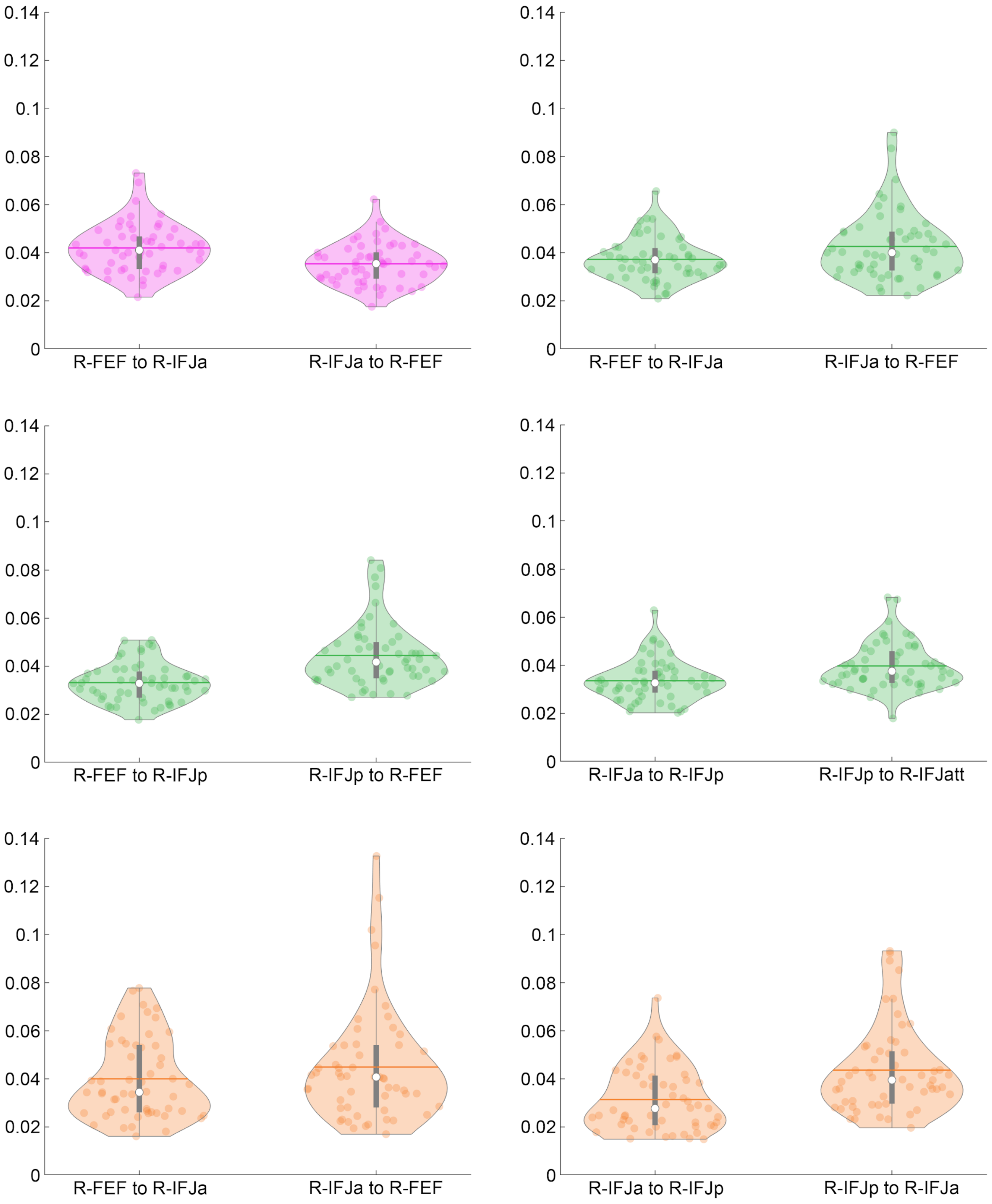


FIGURE S7 The distribution of subjects' PDC value for each significant directional interaction in the right hemisphere. The color of the plots represents different frequency bands illustrated in Figure 5. The dots correspond to each subject's PDC value. The horizontal lines and white dots show the mean and median, respectively.

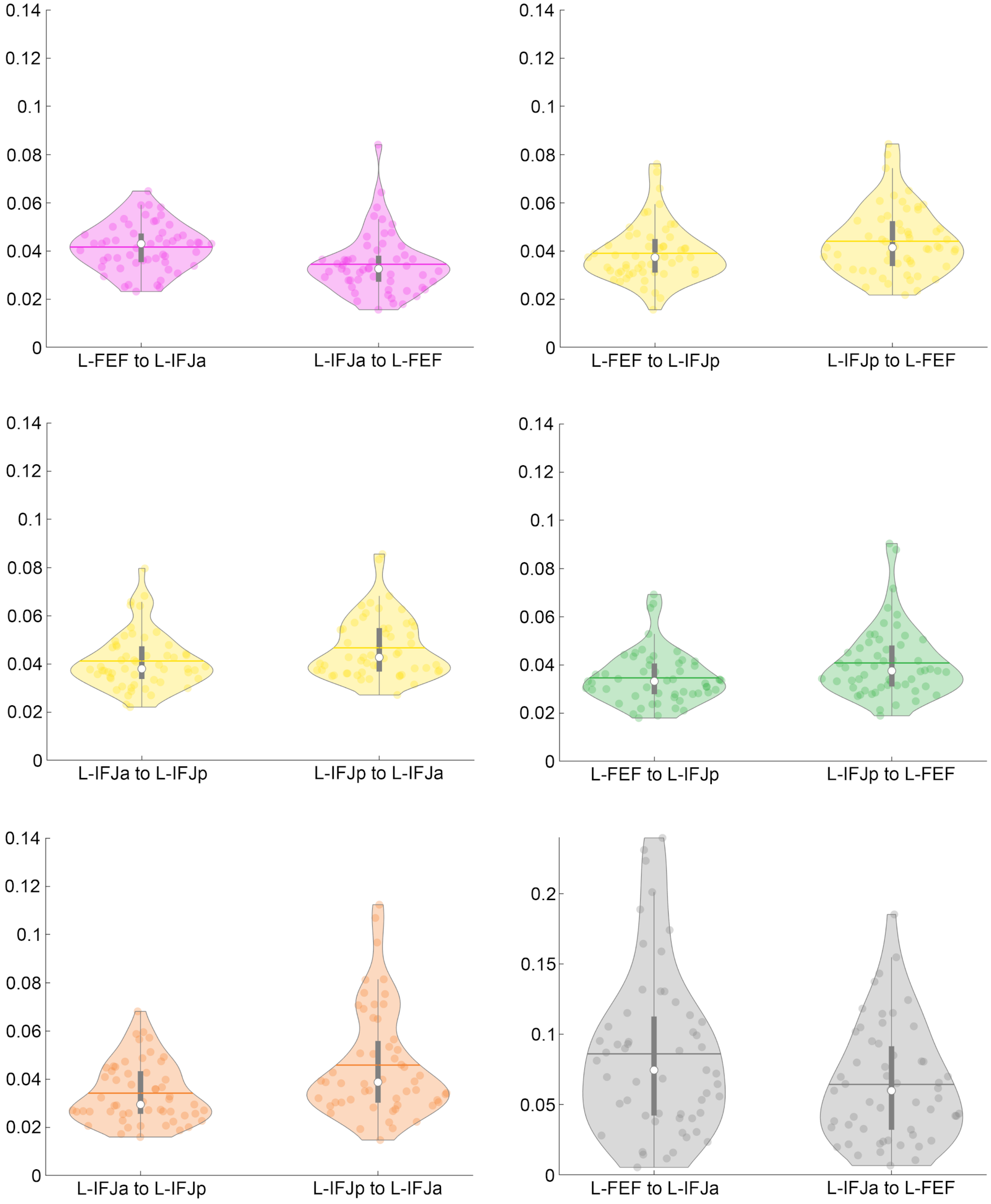


FIGURE S8 The distribution of subjects' PDC value for each significant directional interaction in the left hemisphere. The color of the plots represents different frequency bands illustrated in Figure 5. The dots correspond to each subject's PDC value. The horizontal lines and white dots show the mean and median, respectively.

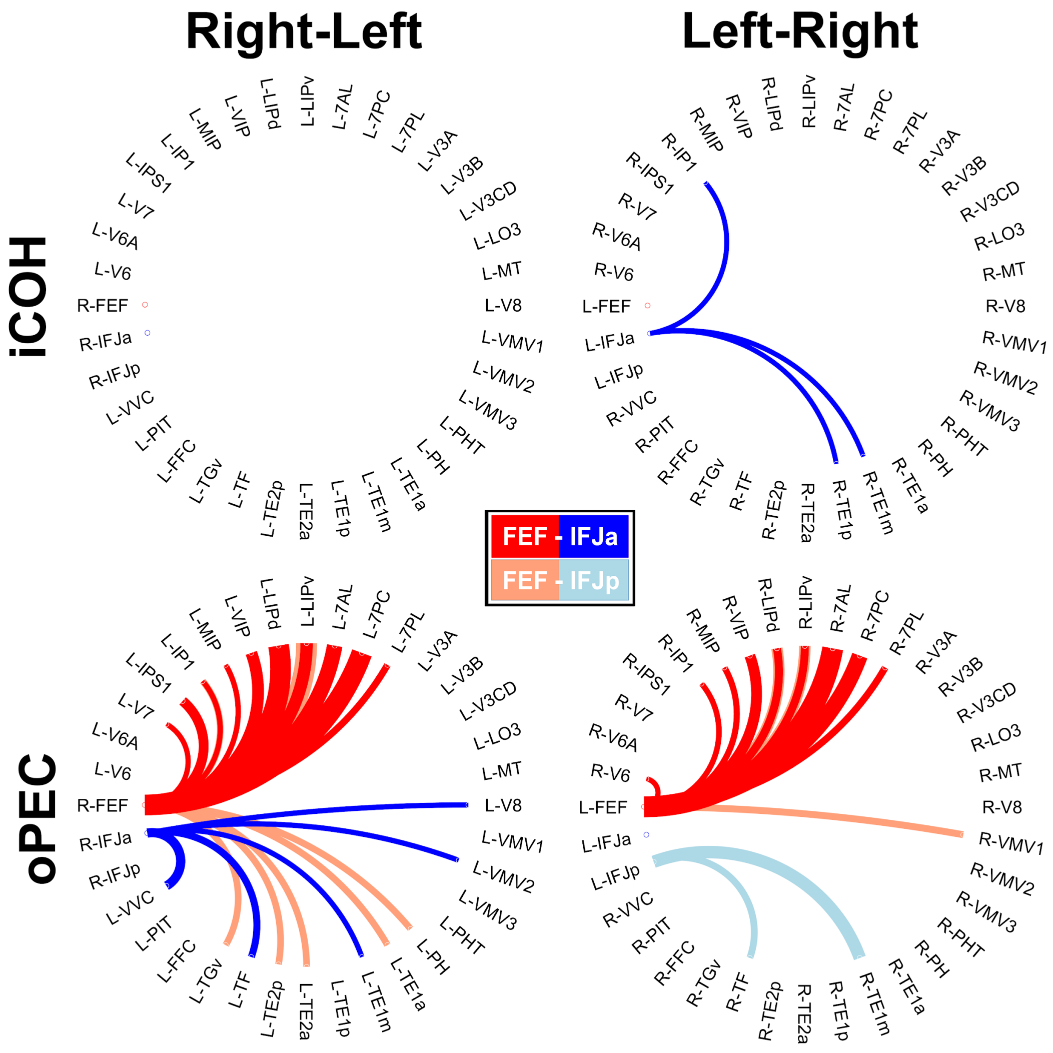


FIGURE S9 Contralateral ROI analyses for the predominant functional connectivities of FEF, IFJa, and IFJp. The width of a line reflects its relative z-score to others (two-sided Wilcoxon signed-rank test, $p < 0.05$, FDR-corrected for 33 ROIs). The results shown here are based on 2 s epoch segmentation.

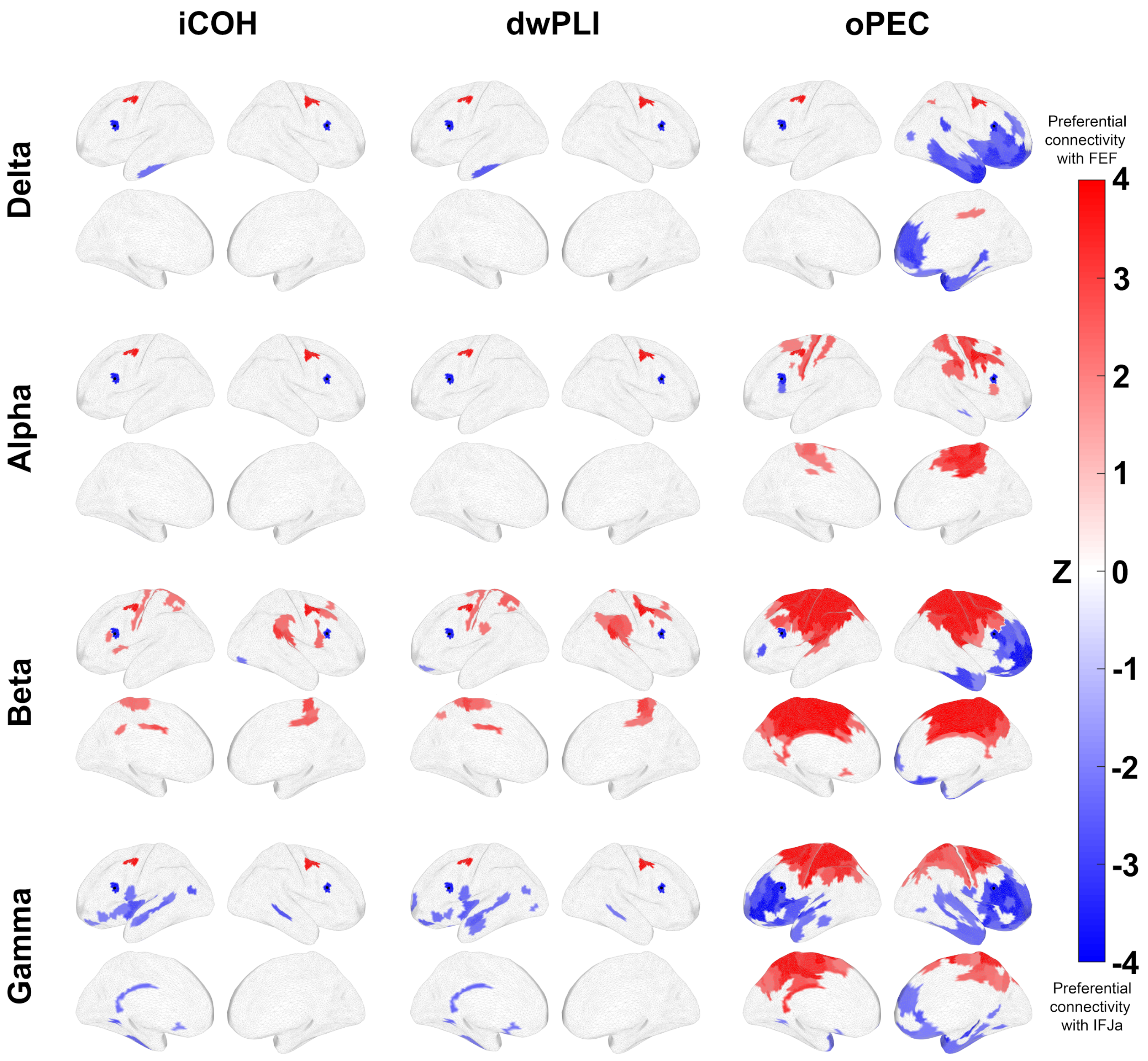


FIGURE S10 Whole-brain exploratory analyses for the predominant functional connectivity profiles of FEF and IFJa (two-sided Wilcoxon signed-rank test, $p < 0.05$, FDR-corrected for 180 parcels). The results shown here are based on 2 s epoch segmentation.

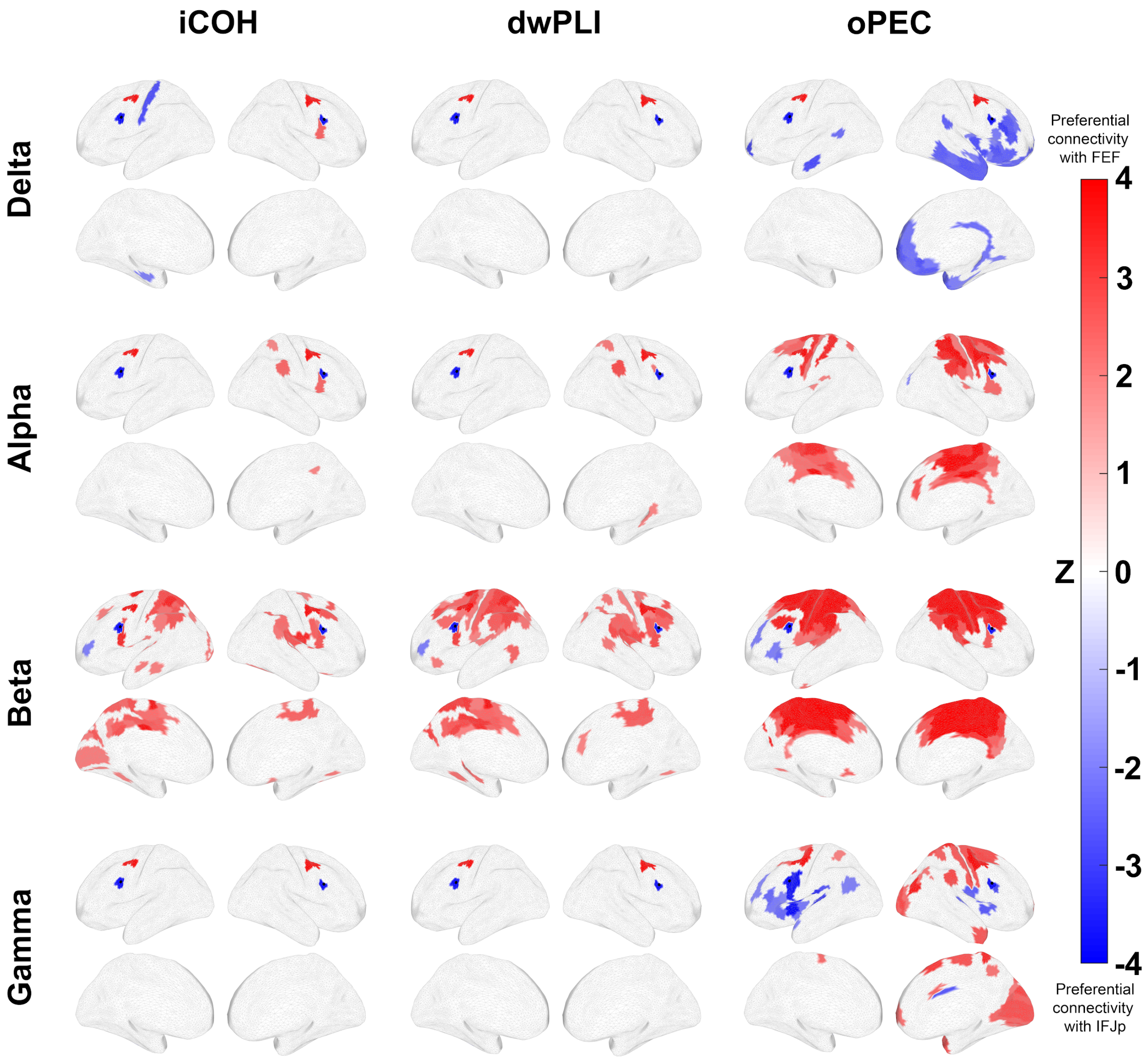


FIGURE S11 Whole-brain exploratory analyses for the predominant functional connectivity profiles of FEF and IFJp (two-sided Wilcoxon signed-rank test, $p < 0.05$, FDR-corrected for 180 parcels). The results shown here are based on 2 s epoch segmentation.

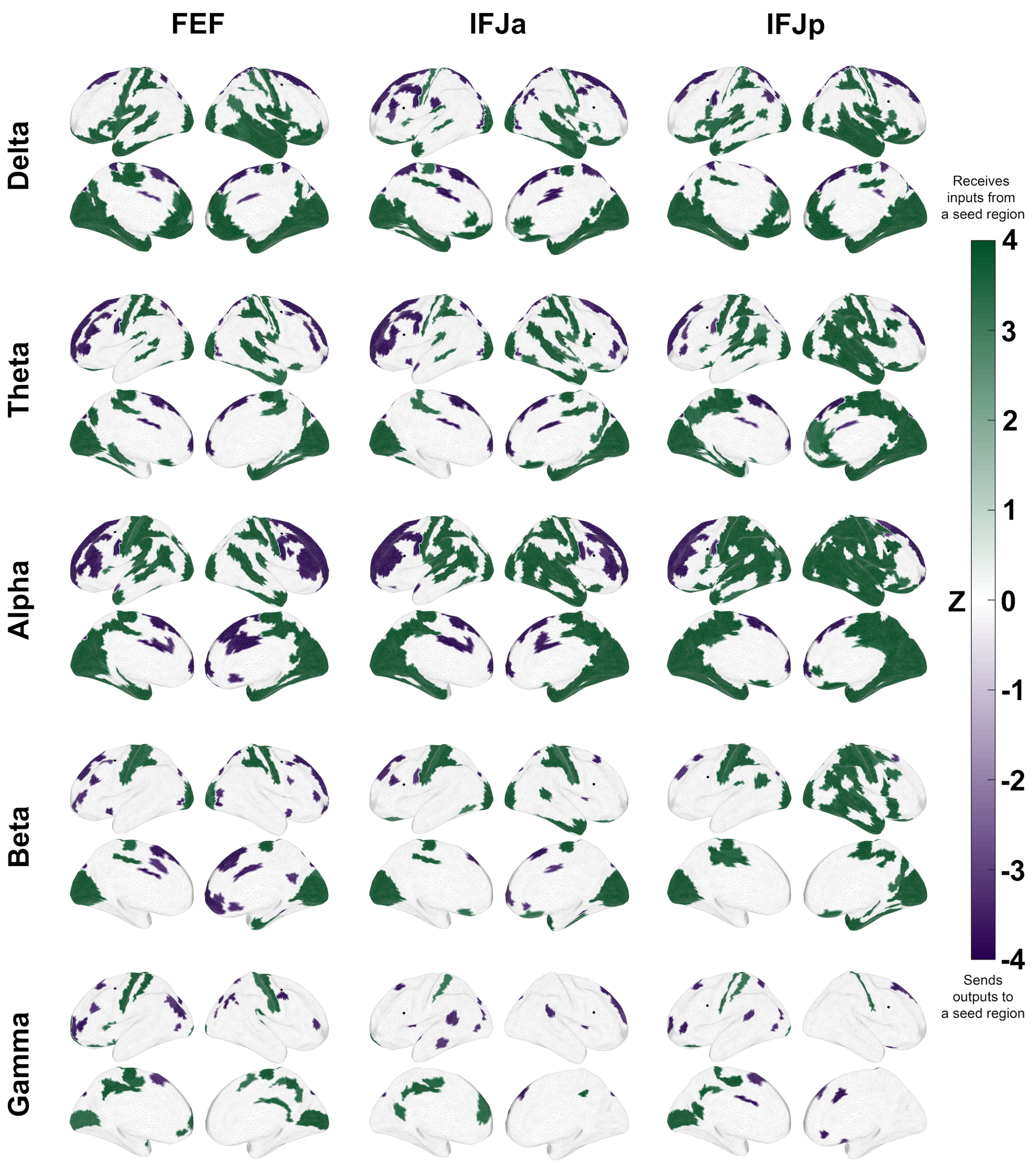


FIGURE S12 Whole-brain exploratory analyses for partial directed coherence. The figure shows the directional influences between FEF, IFJa, IFJp, and the rest of the brain. While the green regions receive inputs, the magenta ones send outputs to a seed region (two-sided Wilcoxon signed-rank test, $p < 0.001$, FDR-corrected for 180 parcels). The results shown here are based on 2 s epoch segmentation.

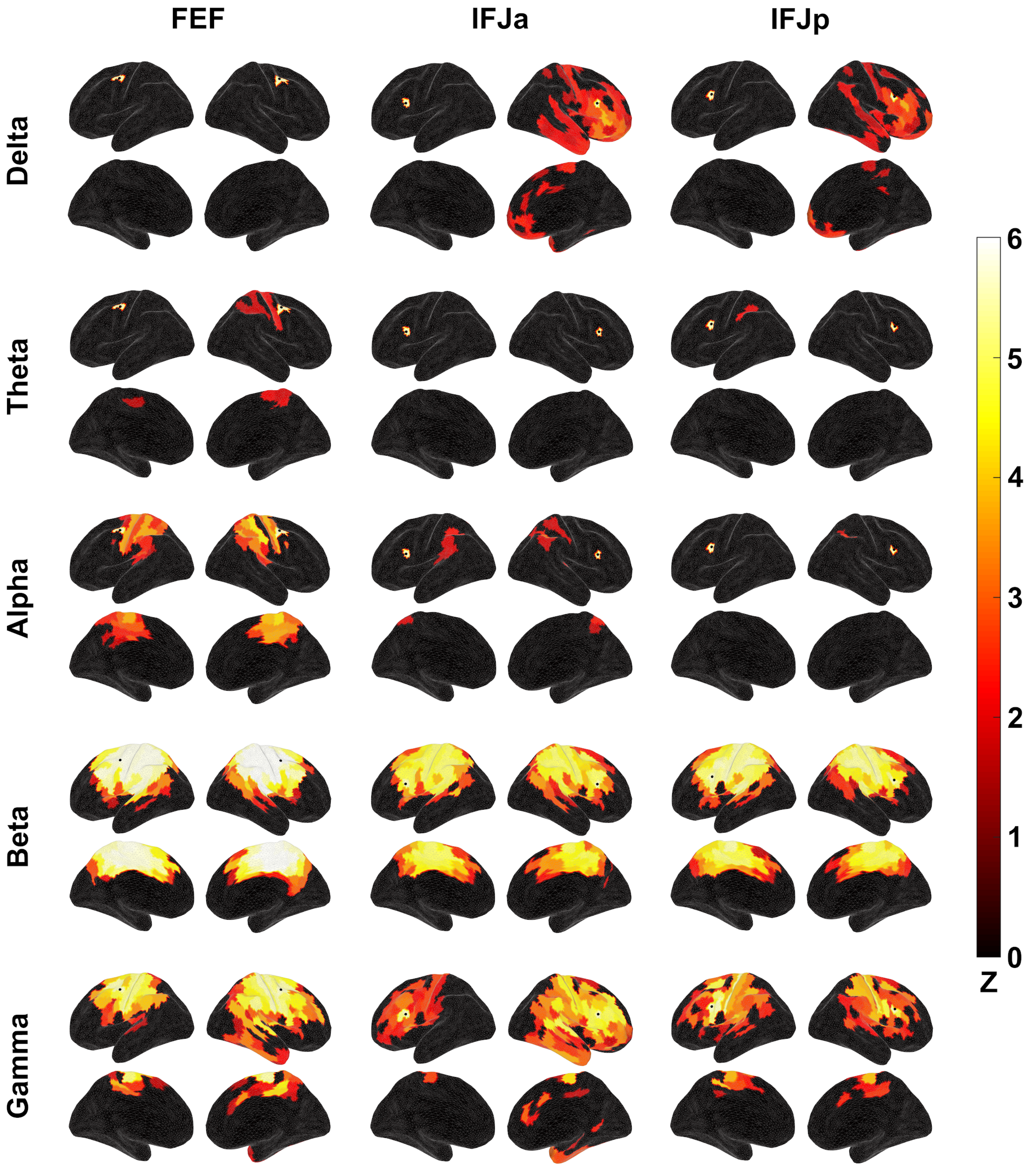


FIGURE S13 The intrinsic power correlation (oPEC metric) of FEF, IFJa, and IFJp with the rest of the brain (one-sided Wilcoxon signed-rank test, $p < 0.05$, FDR-corrected for 180 ROIs) . The results shown here are based on 2 s epoch segmentation.

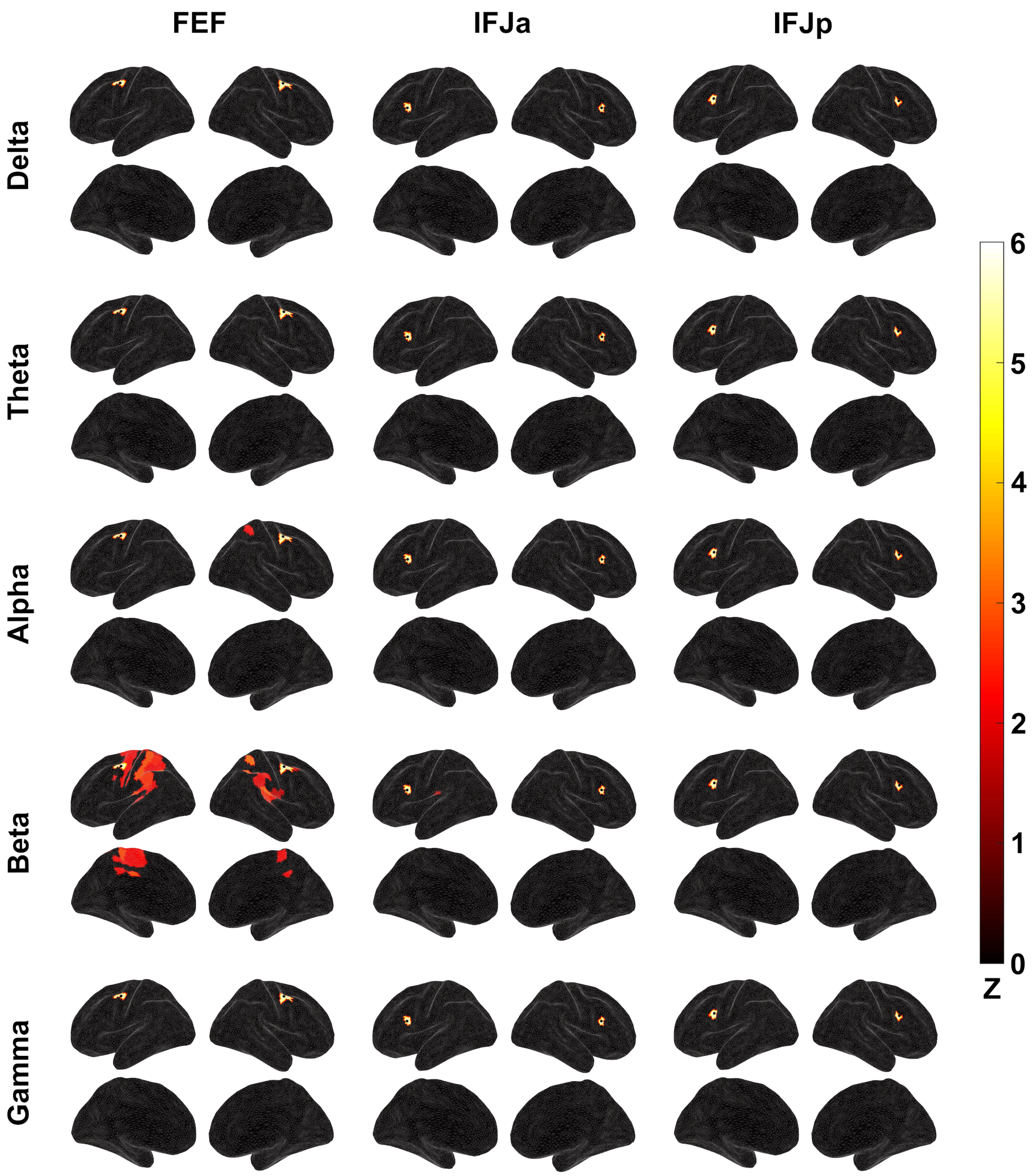


FIGURE S14 The intrinsic power correlation (iCOH metric) of FEF, IFJa, and IFJp with the rest of the brain (one-sided Wilcoxon signed-rank test, $p < 0.05$, FDR-corrected for 180 ROIs) . The results shown here are based on 2 s epoch segmentation.
